# A complete-genome view of phylum Omnitrophota and a multi-order capacity for very long proteins

**DOI:** 10.64898/2026.06.07.730766

**Authors:** Torben N. Nielsen, Lauren M. Lui

## Abstract

Phylum *Omnitrophota* (formerly candidate division OP3) is represented in public databases almost entirely by metagenome-assembled genomes; GTDB R232 contains two complete *Omnitrophota* assemblies. We present 176 complete and 53 high-quality *Omnitrophota* genomes from Oxford Nanopore metagenomes of Fennoscandian deep groundwater and the Baltic Sea water column, an 88-fold expansion of the complete *Omnitrophota* count. The 229 genomes resolve to 202 distinct species at 95 % ANI; 162 of these have no conspecific match in the 714 NCBI HQ *Omnitrophota* MAGs. Phylogenomically, 171 of the 176 complete genomes fall in class *Gorgyraeia* — which contains the cultured episymbiont *Velamenicoccus archaeovorus* — and 5 in *Omnitrophia*, with multiple *Gorgyraeia* orders and families represented. Phylum *Omnitrophota* hosts many very long proteins, with the longest in our corpus reaching 147,155 AA; the long end of the length distribution is concentrated on *Gorgyraeia* contigs across multiple *Gorgyraeia* orders. 24–28 % of GTDB-Tk-classified *Omnitrophota* contigs in the deep-groundwater and Baltic samples host at least one ≥ 10-kAA protein. Across the 916-protein long-protein domain-architecture catalog, 94 % carry transmembrane helices or a signal peptide; the four complete-genome proteins above 100,000 amino acids are all in inner-membrane-anchored architectures, the 147,155-AA protein with 147 TM helices. The 176 complete genomes share a uniform metabolic profile across the dominant orders: intact bacterial peptidoglycan biosynthesis alongside fragmentary TCA, incomplete electron transport, absent aerobic terminal oxidase, and partial cofactor and amino-acid biosynthesis. The profile matches the cultured *V. archaeovorus* phenotype and is consistent with a host-dependent episymbiotic lifestyle. Hypervariable-region calling across the 229 chromosomes returns 1,909 candidate loci, distributed across 223 of them; ribosomal-protein and EF-Tu / EF-G content sits inside called HVRs on 150 of those 223 (67 %), recovering across the collection the housekeeping-cargo integrations documented in Nielsen (2026b). All genomes, the OrthoFinder supermatrix and its ML tree, the 916-protein giant-protein domain-architecture catalog, and per-step scripts are released as a community resource at Zenodo (DOI [TBD]).

## Introduction

The bacterial phylum *Omnitrophota* (formerly candidate division OP3) sits within the PVC superphylum alongside *Planctomycetota, Verrucomicrobiota, Chlamydiota*, and several smaller groups, and is represented in public databases almost entirely by metagenome-assembled genomes (Glöckner et al. 2010; Rinke et al. 2013; Perez-Molphe-Montoya et al. 2022; Seymour et al. 2023). The phylum spans at least six classes in GTDB R232, more than 700 species, and occurs in freshwater, marine, terrestrial-subsurface, and host-associated habitats. The cultured *Gorgyraeia* representative *Velamenicoccus archaeovorus* attaches to and lyses methanogenic archaea in anaerobic enrichment cultures (Kizina et al. 2022). Complete genomes are rare: GTDB R232 holds two — *V. archaeovorus* (GCF_004102945.1) and one *Zapsychrales* assembly (GCA_963662715.1). The remaining ~ 700 species representatives are MAGs of varying completeness or unclosed isolate assemblies.

By *complete genome*, we mean a single contig labeled circular by the assembler with CheckM2 completeness ≥ 90 % and contamination < 5 %, and by *high-quality genome*, a single contig meeting the same CheckM2 cutoffs but not labeled circular. Both definitions are more restrictive than the GTDB high-quality threshold, which admits assemblies fragmented across multiple contigs. Where the circularity distinction does not affect the analysis we combine the two tiers.

Most bacterial proteins are short: the typical bacterial proteome has fewer than 1 % of its proteins above 1,000 AA, and proteins above 5,000 AA are rare across the tree of life and concentrated in a small number of bacterial phyla (Reva & Tümmler 2008; Amaral & Devos 2024). The largest known bacterial proteins are in the multi-kAA range — *Salmonella* SiiE at 5,559 AA, *Mycobacterium ulcerans* mlsA1 at ~17 kAA, *Chlorobium chlorochromatii* CaD3’s 36,806 AA parallel-β-helix protein — and these top out an order of magnitude below the longest *Omnitrophota* proteins previously reported (up to 85,804 AA; West-Roberts et al. 2023). Whether multi-kAA proteins are a property of *Omnitrophota* as a phylum or an artifact of the small number of available assemblies could not be tested without a substantially larger complete-genome collection.

This paper uses Oxford Nanopore long-read metagenomes from two Fennoscandian deep-groundwater sites (KR0015B at 69 m below sea level; SA1420A at 201 m below sea level, both at the Äspö Hard Rock Laboratory in Sweden) and 16 Baltic Sea water-column samples to address that question at collection scale. We detect GTDB-Tk-classified *Omnitrophota* contigs at both Fennoscandian sites and in three Baltic samples (B5, B7, B10). From these we recover 176 complete and 53 high-quality *Omnitrophota* genomes, an 88-fold expansion over the two prior complete *Omnitrophota* assemblies.

We use the collection for three analyses. First, we place the new genomes phylogenomically within *Omnitrophota* at class and order. Second, we document a giant-protein phenotype concentrated in class *Gorgyraeia*: 14 distinct proteins ≥ 80 kAA all on *Gorgyraeia* contigs, plus enrichment at the ≥ 10-kAA threshold across the *Omnitrophota* orders sampled. Third, we characterize chromosome-level hypervariable and mobile-element loci across 223 of 229 chromosomes and find that ribosomal-protein operons appear inside integration/HVR windows on 150 of those 223 — putting a collection-wide rate on Nielsen (2026b)’s finding of a ribosomal-protein operon carried on an integrative mobile element.

## Results

### 229 complete and high-quality *Omnitrophota* genomes from five samples

Of the 18 *Omnitrophota*-candidate samples surveyed (2 Fennoscandian deep-groundwater + 16 Baltic Sea water-column), 5 yielded GTDB-Tk-classified *Omnitrophota* contigs (KR0015B, SA1420A, B5, B7, B10), and each contributed at least one genome to the final inventory. The collection comprises 229 *Omnitrophota* genomes: 176 complete and 53 high-quality (definitions per Methods §”Quality and topology”). The 229 resolve to 202 distinct species at 95 % ANI (skani; 160 species among the 176 complete genomes, 52 among the 53 high-quality, with 10 clusters spanning both tiers); 162 of the 202 species have no conspecific match in the 714 NCBI HQ *Omnitrophota* MAGs. Per-sample provenance of the 176 complete genomes: 77 from KR0015B and 93 from SA1420A (both Fennoscandian deep groundwater), plus 5 from Baltic B10 and 1 from Baltic B7. The 53 high-quality genomes split as 32 KR0015B, 17 SA1420A, 1 B5, 2 B7, and 1 B10 (Fig. 1).

**Figure 1.**
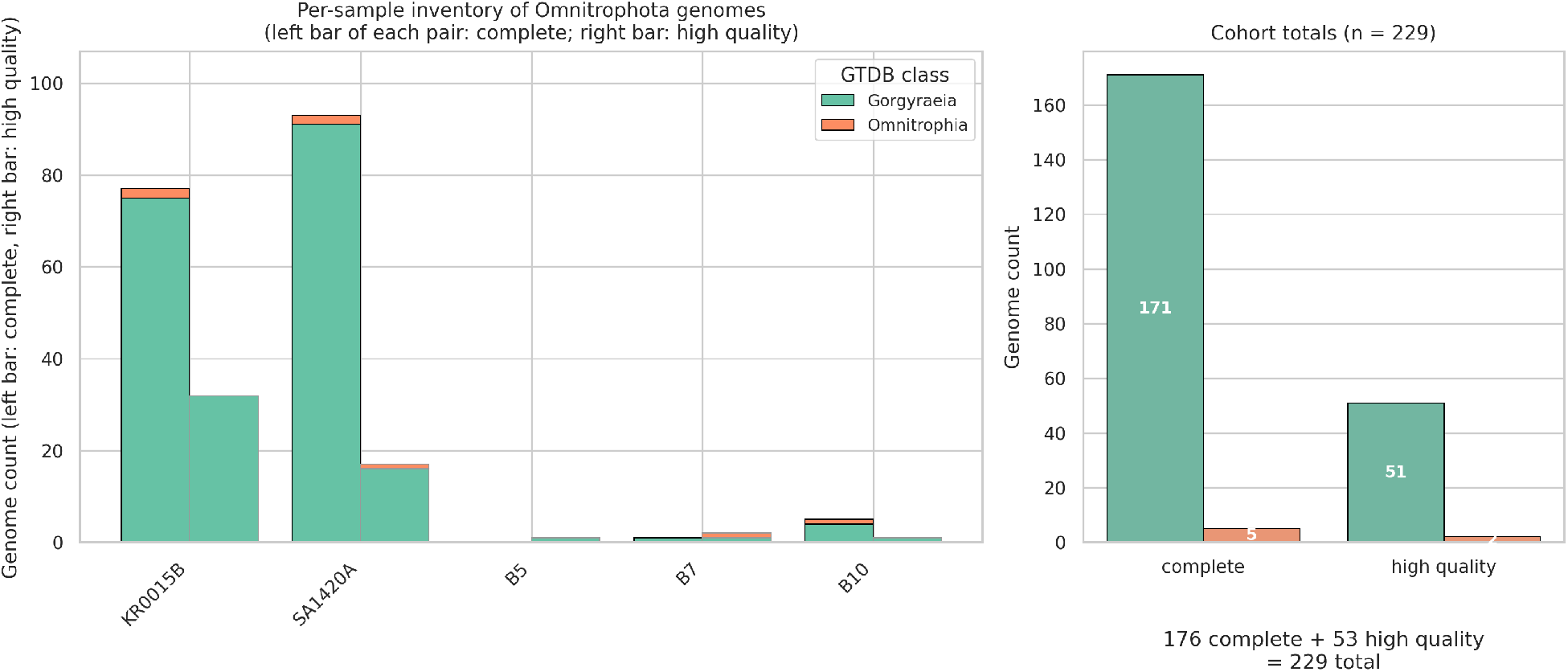
Genome inventory and provenance. 176 complete + 53 high-quality across 5 of 18 samples that yielded any *Omnitrophota* (KR0015B, SA1420A, B5, B7, B10); per-sample stacked bars at GTDB-Tk class resolution. Per-order detail is reported in Supplementary Table S1.

GTDB-Tk R232 assigned the 176 complete genomes across 22 orders and 28 families within phylum *Omnitrophota* — 20 of the 41 *Gorgyraeia* orders in GTDB R232 (49 %) plus both *Omnitrophia* orders, recovered from two Fennoscandian deep-groundwater samples. The largest single order is *Gygaellales* (64 genomes: 63 in family *Profunditerraquicolaceae*, 1 in family *JAHJCI01*), followed by order 2-01-FULL-45-10 (16, family 2-01-FULL-45-10), *Tantalellales* (14, *Tantalellaceae*), and *Gorgyraeales* (14, *Gorgyraeaceae*) within class *Gorgyraeia*. Full per-order counts are reported in Supplementary Table S1.

### Phylogenomic structure

A supermatrix of 97 single-copy orthogroups derived from OrthoFinder on a 184-proteome input set was aligned with MAFFT L-INS-i, trimmed with BMGE v2.0, and inferred under IQ-TREE v3.0.1 with the LG+F+R10 model and 1,000 ultrafast bootstrap replicates (5 h 32 min wall-clock on 32 threads). The input set was passed to OrthoFinder in a single all-vs-all run, so the 97 single-copy orthogroups are the markers conserved across *every* proteome in the input — including the 4 NCBI references and the 4 PVC outgroups.

The 4 NCBI proteomes were included not as additional data but as named taxonomic anchors so that clade labels on the collection tree can be read off the named neighbour rather than inferred genome-by-genome from GTDB-Tk: the cultured *Gorgyraeia* episymbiont *Velamenicoccus archaeovorus* (GCF_004102945.1) and the *Zapsychrales* complete assembly (GCA_963662715.1) — the two prior complete *Omnitrophota* in GTDB R232 — plus two GTDB *Gorgyraeia* species representatives that are HQ MAGs (GCA_020341815.1, GCA_020346285.1). Their participation in OG discovery is a deliberate tradeoff: the marker set is more conservative than a 176-only run would be, but the resulting tree carries named class- and order-level anchors that label our collection clades directly. The 4 anchors do not contribute to phylum-level claims. The set was completed with 4 sample-derived PVC outgroup proteomes — drawn from in-house genomes we had on hand from other work in the project — classified by GTDB-Tk as *Verrucomicrobiia* (n = 2), *Planctomycetia* (n = 1), and *Phycisphaerae* (n = 1), used to root the tree.

Of the 176 complete collection genomes, 171 fall in class *Gorgyraeia* and 5 in class *Omnitrophia* (GTDB-Tk R232 assignments); the collection spans multiple *Gorgyraeia* orders including the giant-protein-bearing *Gygaellales, Duberdicusellales*, and *Gorgyraeales* (Fig. 2). A larger 938-tip supermatrix tree placing the new genomes in the broader GTDB R232 *Omnitrophota* MAG context was computed via the cohort-HMM recruitment pipeline of Nielsen (2026a) and is shown in Supplementary Figure S2; its main-text role is taxonomic placement only, and none of this paper’s headline claims depend on its topology.

**Figure 2.**
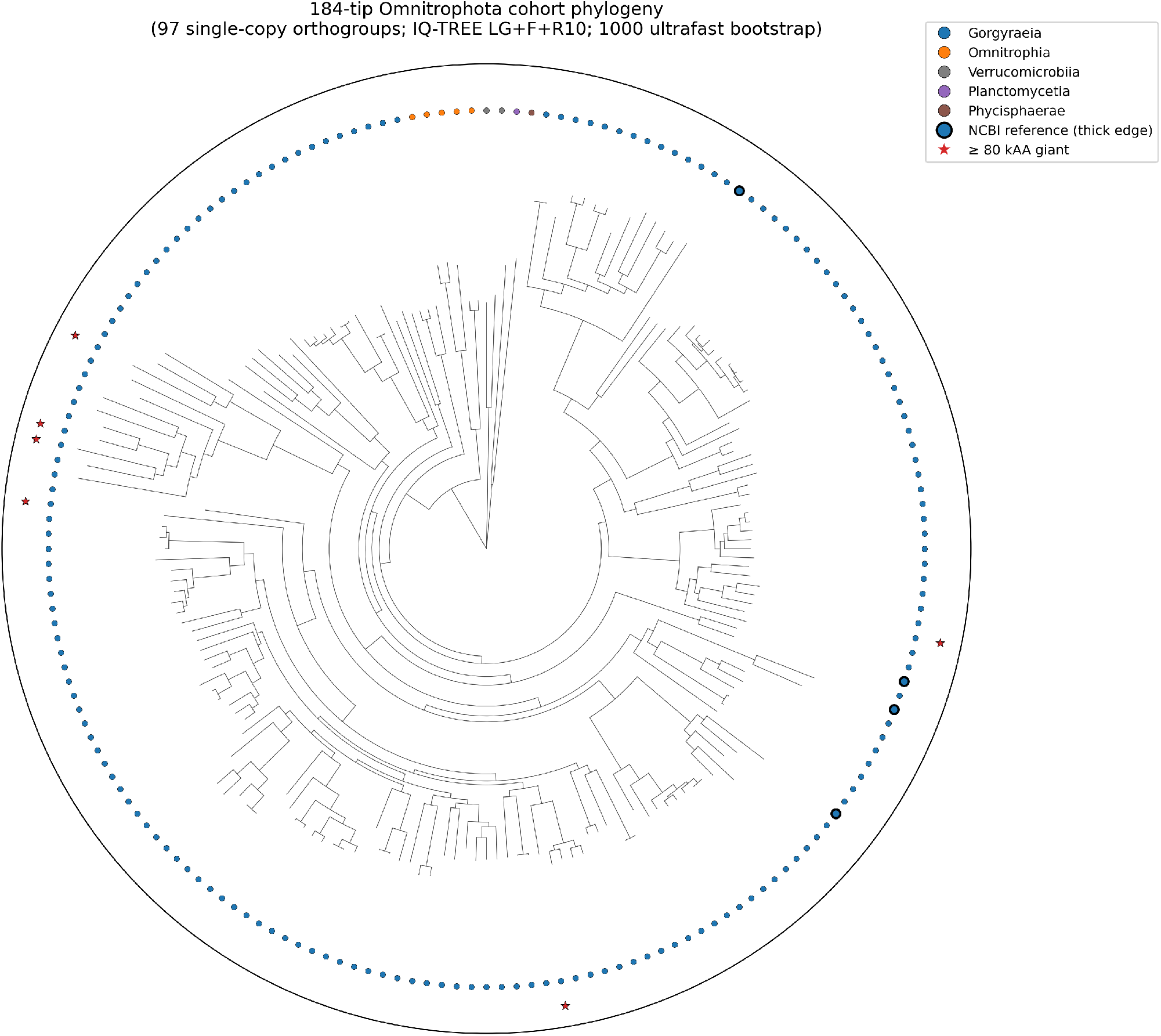
Collection phylogeny. 184-tip OrthoFinder 97-SC-OG supermatrix IQ-TREE radial layout (`phylogeny/plot_radial_tree_184.R`). Tips colored by GTDB-Tk class (*Gorgyraeia* dominant, *Omnitrophia* minor) and by sample (KR0015B / SA1420A / Baltic); giant-protein-bearing genomes marked; outgroup PVC tips collapsed for clarity.

### A multi-order capacity for very long proteins

We scanned Pyrodigal protein-coding predictions across the 18 *Omnitrophota*-candidate samples — the 2 Fennoscandian deep groundwater sites and the 16 Baltic Sea water-column samples (B1–B12, B14, B15, B17, B19) — for predicted proteins of unusual length. The long-protein landscape is concentrated in *Omnitrophota* (Fig. 3):

**Figure 3.**
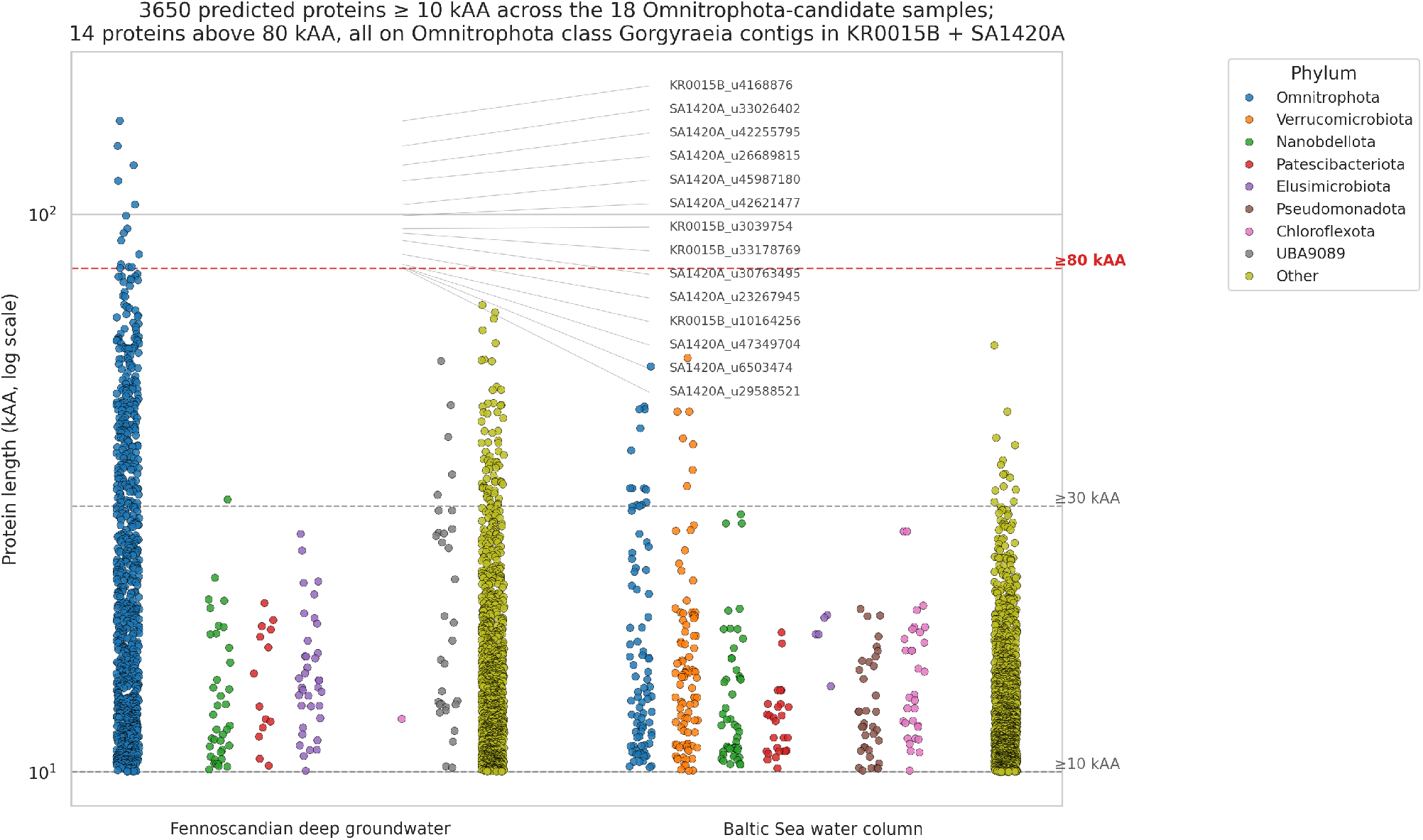
Very long proteins are concentrated in Omnitrophota. Protein-length distribution (≥ 10 kAA, log scale) as a stripchart across the two sample groups, split by phylum. The other phyla are shown as a baseline: long proteins occur across the community, but every protein above 80 kAA — 14 in total, called out individually with sample and order labels — is on an Omnitrophota (class *Gorgyraeia*) contig in the two Fennoscandian deep-groundwater samples KR0015B and SA1420A.

- 3,650 proteins ≥ 10 kAA, 348 ≥ 30 kAA, 97 ≥ 50 kAA, and 14 ≥ 80 kAA across the corpus.
- The long end of the corpus-wide length distribution is *Gorgyraeia*-concentrated. The 14 longest proteins — all above 80 kAA — come from the two Fennoscandian deep groundwater samples (KR0015B and SA1420A), and each one sits on a contig classified by GTDB-Tk as *Omnitrophota* class *Gorgyraeia* — the class containing the cultured episymbiont *Velamenicoccus archaeovorus*. They are 14 distinct proteins on 14 distinct contigs: an all-vs-all DIAMOND blastp of the set yields zero pairs above 30 % identity over more than 30 % of the shorter member, so no two of the 14 are pairwise similar at this 30 % / 30 % threshold and they are sequence-level distinct. One cross-member alignment runs the full length: a 122,534-AA and a 99,464-AA *Gorgyraeaceae* giant (`SA1420A_u42255795_1875` and `SA1420A_u42621477_744`) match at 58.6 % identity over essentially the full length. This single cross-pair indicates that the giant-protein architecture can be conserved across divergent members of the same family rather than being a strain-specific feature of one circular genome, but does not on its own demonstrate family-wide conservation.
- Of the 14 contigs hosting these proteins, six are circular and are members of our 229-genome collection; eight are non-collection *Omnitrophota* contigs of 0.78–3.76 Mbp that fall below the collection’s CheckM2 thresholds. The cross-sample protein-length scan operates at per-sample Pyrodigal output and therefore reports all 14 regardless of collection membership; restricting to the 229 would drop 8 *Gorgyraeia* giants observed on independent contigs, none of which are highly similar to the 6 collection giants by all-vs-all DIAMOND.
- Five of the eight excluded contigs were excluded on the contamination side — CheckM2 contamination 8.4–23.6 % at completeness ≥ 91 % — rather than on completeness. Each is a *single* 2–3 Mbp contig, so conventional bin-level cross-contamination is not the mechanism. Two non-exclusive mechanisms are consistent with the contamination signal: (i) the multi-kAA giant proteins themselves carry duplicated domain content that CheckM2 reads as duplicated single-copy markers, and (ii) collapsed multi-copy assembly inflates marker counts directly. Three of the five excluded contigs carry myloasm-reported Multiplicity > 1.0 (1.03–1.11×), supporting (ii) for those three. We have not run the direct CheckM2-marker-versus-giant-protein-CDS overlap test that would discriminate (i) from (ii); both should be treated as candidate explanations rather than established mechanism.
- The remaining three excluded contigs failed on completeness (29.6 %, 40.1 %, 70.8 %); two of these three (`KR0015B_u10164256`, `SA1420A_u42621477`) are also the only two excluded contigs at < 15× coverage. The exclusion pattern is therefore not a quality-of-protein-call signal.
- The longest predicted protein in the entire corpus is 147,155 AA, on KR0015B_u4168876 (*Omnitrophota* class *Gorgyraeia*, order *Gygaellales*, family *Profunditerraquicolaceae*).
- The 20 longest predicted proteins across the corpus are likewise 100 % *Omnitrophota* class *Gorgyraeia*.

Taxonomic breakdown of long proteins by sample group (Table 1):

| Sample group | $\geq 10$ kAA n | $\geq 30$ kAA n | $\geq 80$ kAA n | $\geq 30$ kAA dominant phyla |
| --- | --- | --- | --- | --- |
| Fennoscandian deep groundwater (KR0015B + SA1420A) | 2,108 | 314 | 14 | *Omnitrophota* 240/314 (76%) |
| Baltic Sea water column (16 samples) | 1,542 | 34 | 0 | *Omnitrophota* 13, *Verrucomicrobiota* 7, others 14 |

The *Omnitrophota* density of long proteins is not a denominator artifact. We counted GTDB-Tk-classified *Omnitrophota* contigs per sample group: 1,679 in KR0015B+SA1420A and 144 in the Baltic Sea. These denominators are sample-wide — every classified contig regardless of CheckM2 quality, not restricted to the 229-genome collection. Among classified *Omnitrophota* contigs, 28.0 % in Fennoscandian deep groundwater and 24.3 % in the Baltic water column host at least one ≥ 10-kAA protein, and 13.4 % and 6.2 % respectively host at least one ≥ 30-kAA protein.

No *Verrucomicrobiota* protein reaches 80 kAA in either sample group, so the giant-tier phenotype is *Omnitrophota*-specific. At the more permissive ≥ 30-kAA threshold, the Baltic samples do show a possible PVC-wide extension — 7 *Verrucomicrobiota* alongside 13 *Omnitrophota* — but at *n* = 7 this stays suggestive rather than established.

The long-protein signal is concentrated in the complete tier and depleted in NCBI HQ MAGs at the same taxonomic scope (Fig. S3). Across the 714 NCBI HQ *Omnitrophota* MAG population — GTDB R232 species representatives at CheckM2 completeness ≥ 90 % and contamination < 5 % — 306, or 42.9 %, host at least one ≥ 10-kAA protein. In our 176 complete genomes, 165 (93.8 %) host at least one, with a median of 2 long proteins per genome versus 0 for the NCBI population. The 53 in-house HQ-tier genomes sit between at 81.1 %, indicating that assembly tier contributes to the gap independently of provenance; a residual provenance or pipeline effect on top of that cannot be ruled out from these data.

The driver is assembly contiguity, not gene-content completeness. Long-protein count across these 714 NCBI MAGs is anti-correlated with contig count, Spearman ρ = −0.36, p < 1 × 10^−23^, and the response is monotonic: 100 % of 1–5-contig MAGs carry at least one ≥ 10-kAA protein versus 17.9 % at 201–500 contigs. A ≥ 10-kAA ORF needs roughly 30 kb of intact coding sequence on a single contig; MAGs with N50 in the tens of kb break long ORFs at contig boundaries, where Pyrodigal calls them as two truncated proteins instead of one. CheckM2’s KO-presence-based score does not penalize this fragmentation. The HQ threshold therefore admits MAGs that are systematically incapable of resolving the long-protein landscape — a resolution that requires complete-tier assembly.

### Three architectural classes of giant protein

InterProScan (IPS) was run against an architectural member-database panel — Pfam, SMART, CDD, PANTHER, ProSitePatterns, ProSiteProfiles, Gene3D, Coils, and MobiDBLite, with SUPERFAMILY excluded for the technical reason given in Methods — on every *Omnitrophota* protein ≥ 10 kAA in the collection. The full set comprises 912 main-run proteins ≤ 99,999 AA from the 229-genome collection and the four collection ≥ 100-kAA proteins processed via a chunked-rescan and canonical-zone stitching pipeline (`scripts/27_chunk_giants_rescan.py`, `scripts/28_giants_stitch_results.py`) that works around the hard 100,000-AA per-target ceiling inside HMMER 3.x’s pipeline. The 8 ≥ 80-kAA giants on non-collection *Omnitrophota* contigs were run separately (`scripts/27b_chunk_giants_noncirc.py`, `scripts/ 28b_giants_stitch_noncirc.py`, `scripts/49_combine_ips_all14.py`) so that the all-14-giant analysis is on identical IPS member-database settings. 779 of 916 long proteins (85 %) carry at least one Pfam hit; the remaining 137 are Pfam-dark-matter, with no hits across any of the nine InterProScan member databases listed above (Fig. 4).

**Figure 4.**
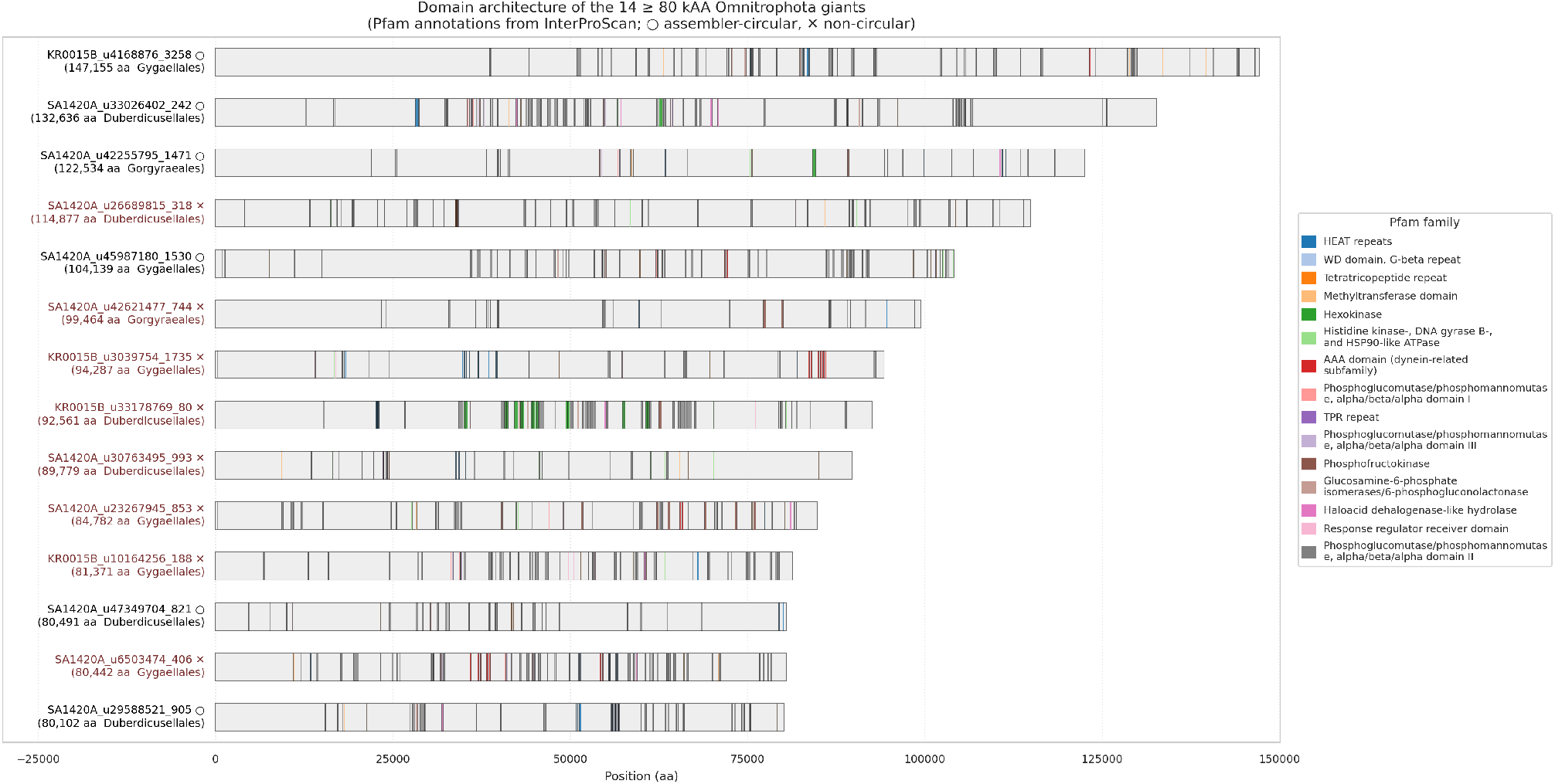
Giant-protein domain architecture. Linear domain cartoons for all 14 ≥ 80-kAA giants (6 collection + 8 non-collection) drawn to scale, arranged by decreasing length from the 147,155-AA example at top to the shortest near 80 kAA at the bottom (within the 80–99-kAA band the ordering is approximate, not strictly monotone). Pfam annotations from InterProScan; the 779 of 916 long proteins (85 %) with at least one Pfam hit and the 504 / 779 (65 %) with at least one Pfam repeated ≥ 2× in the same polypeptide are summarized in §”Three architectural classes”. Top-15 most-recurring Pfam families colored.

Across the 779 Pfam-positive long proteins, 552 distinct Pfam families occur. A Pfam co-occurrence graph at the protein level (`scripts/26_ips_domain_cooccurrence.py`) with edge weight equal to the number of proteins jointly carrying both Pfams, restricted to count ≥ 3 and positive pointwise mutual information (PMI), partitions into 10 greedy-modularity communities of recognizable functional content. The two largest modules each cover several hundred long proteins. Module 1 is centered on signal-transduction / regulatory architecture — methyltransferase domain (PF08241, 137 proteins), histidine-kinase / HSP90-like ATPase (PF02518, 136), GNAT acetyltransferase (PF00583, 93), HAD hydrolases (PF13419, 74), and HD-domain regulators. Module 2 is centered on sugar metabolism — starch synthase catalytic (PF08323, 153), glycosyl transferase group 1 (PF00534, 127), phosphoglucomutase / phosphomannomutase (PF02878 / PF02879 / PF02880; 88 / 65 / 64), response-regulator receiver (PF00072, 105), radical SAM (PF04055, 90), and ROK-family kinase (PF00480, 44). The remaining modules carry, in turn, transport / membrane architecture, oxidoreductase content, peptidase / protease activity, repeat scaffolding (HEAT / NHL / TPR), and smaller specialized clusters.

Three architectural classes emerge from the per-protein repeat structure (`scripts/ 30_long_protein_repeats.py`):

1. **Multi-enzyme metabolic clusters**. Many distinct Pfams tile the polypeptide with no single Pfam dominating by copy number. The longest example is `KR0015B_u4168876_114` at 147,155 AA, which carries 43 distinct Pfams spanning a sugar-metabolism cluster, a kinase / phosphotransferase cluster, and scattered methyltransferase, NUDIX, HIT, signal-peptidase-II, and F420-oxidoreductase content with internal HEAT and Adaptin-N scaffolding.
2. **Tandem-repeat scaffolds**. One Pfam dominates by copy number, embedded in a long polypeptide whose remaining content varies. The most extreme example is a 62,866-AA protein on NCBI assembly GCA_041666865.1 contig JBAYGD010000001.1 with 41 copies of the bacterial TSP3 repeat (PF18884). Other class-2 carriers include high-copy concanavalin-A-like lectin / glucanase (PF13385, up to 36 copies), NHL repeat (PF01436, up to 37 copies), and passenger-associated transport repeat (PF12951, up to 38 copies).
3. **Dockerin-bearing**. Dockerin type I (PF00404) appears in 21 proteins, with up to 15 copies in a single protein. This is the “Archaeovorin” / “Dockerin-type” architecture (West-Roberts et al. 2023) — present in our collection but neither dominant nor exclusive.

Among the 14 ≥ 80-kAA giants specifically, classes 1 and 2 are both well-represented while class 3 is absent. Nine giants are multi-enzyme metabolic — KR0015B_u4168876_114 (147 kAA, 43 distinct Pfams), SA1420A_u33026402_205 (132 kAA, 50 distinct), SA1420A_u42255795_1875 (122 kAA, 27 distinct), SA1420A_u45987180_2000 (104 kAA, 44 distinct), SA1420A_u47349704_1584 (80 kAA, 25 distinct), and four giants on non-collection contigs (KR0015B_u10164256_188 47 distinct; SA1420A_u26689815_318 45 distinct; SA1420A_u23267945_853 35 distinct; SA1420A_u42621477_744 21 distinct). Five giants are tandem-repeat scaffolds — SA1420A_u29588521_868 (21 copies of WD repeat PF00400), SA1420A_u6503474_406 (20 copies of WD repeat), KR0015B_u3039754_1735 (13 copies of HEAT repeat PF13646), KR0015B_u33178769_80 (9 copies of hexokinase PF03727 in a multi-Pfam background), and SA1420A_u30763495_993 (7 copies of TPR PF13414). The eight non-collection giants distribute across the same two classes as the six collection giants (4 / 4 multi-enzyme vs tandem-repeat split among the non-collection contigs; 5 / 1 split among the collection), consistent with the architectural diversity not being a collection-membership artifact, though at *n* = 6 / 8 this is not a test that would reject the null.

504 of 779 Pfam-positive long proteins (65 %) carry at least one Pfam repeated ≥ 2× in the same polypeptide; 114 (15 %) ≥ 5×; 47 (6 %) ≥ 10×; 14 (2 %) ≥ 20×. The repeat phenotype is therefore the rule, not an outlier feature of a handful of cases.

Whether these architectural repeats arose by recent tandem-nucleotide expansion or by ancient duplication followed by sequence divergence is testable directly: we scanned the underlying CDS for tandem nucleotide repeats with both a strict back-to-back identical-copy detector (`scripts/ 34_long_gene_nt_repeats.py`) and a fuzzy Tandem-Repeats-Finder scan that allows mismatches and indels between copies (`scripts/35_long_gene_nt_repeats_fuzzy.py`). Under the strict criterion only 17 % of long genes carry any tandem and 0 % reach 10 % of CDS in tandems; under the fuzzy criterion 86 % carry at least one tandem, 24 % reach 10 % of CDS in tandems, and 5 % reach 50 %. The 17 % that retain strict tandems do so over small fractions of CDS — none above 10 % — and are most likely short, locally stable tandems rather than evidence of recent large-scale expansion. Synonymous and tolerated nonsynonymous substitutions have accumulated between copies, so the nucleotide tandems have dissolved while the protein-domain architecture persists. The repeat architecture is therefore not the product of recent tandem expansion across the bulk of the collection.

Two complementary clustering passes characterize the long-protein corpus at the family level (`scripts/61_long_protein_mmseqs_cluster.py`, `scripts/ 62_long_protein_pfam_architecture_cluster.py`). Sequence-level mmseqs2 easy-cluster across all 5,761 ≥ 10-kAA proteins, in-house and NCBI HQ combined, returns 3,284 clusters at 30 % identity / 80 % coverage of which 2,624 (80 %) are singletons; only 3 clusters exceed 50 members and only 16 span both in-house and NCBI populations — the long-protein landscape is overwhelmingly composed of sequence-unique proteins, with the cross-source mixing further depressed by the coverage filter penalizing NCBI MAGs whose long ORFs are contig-boundary-truncated (Fig. S3). Independent Pfam-architecture clustering on the 779 Pfam-positive proteins at Jaccard 0.7 yields 416 architecture clusters: 281 singletons, 21 medium clusters of 5–19 proteins, and 2 large clusters of 20–49 — a substantially flatter distribution than sequence clustering, consistent with conserved domain architecture against a backdrop of divergent sequence. The two largest architecture clusters are a 24-protein PF03412 (Peptidase C39 family) group and a 20-protein PF07517 (SecA DEAD-like domain) group; the 17-protein PF00404 dockerin + PF18884 TSP3-repeat cluster recovers the “Archaeovorin” architecture (West-Roberts et al. 2023) as a cluster in our data.

### Membrane topology of the giant proteins

DeepTMHMM v1.0 (Hallgren et al. 2022) was run on all 916 long proteins in the domain-architecture catalog to assign each a canonical topology class and a per-residue inside / outside / transmembrane-helix (TM) / signal-peptide (SP) path. The 912 proteins ≤ 99,999 AA were run directly; the 4 collection ≥ 100-kAA giants — including the 147,155-AA protein — sit above DeepTMHMM’s per-target length ceiling and were chunked into ≤ 80-kAA overlapping pieces, predicted per-chunk, and stitched back into full-length per-residue tracks with the same canonical-zone rule used for the InterProScan chunked rescan (`scripts/54_chunk_giants_deeptmhmm.py`, `scripts/ 55_stitch_giants_deeptmhmm.py`; signal-peptide calls suppressed on non-first chunks since an internal ‘S’ on a later chunk is a chunking artifact). Of the 916 proteins screened, 857 (93.6 %) carry at least one transmembrane segment or a signal peptide and only 59 (6.4 %) are predicted globular. Topology class breakdown (Fig. 5): 422 (46.1 %) α-helical TM with no signal peptide, 356 (38.9 %) SP+TM (signal peptide plus TM helices), 79 (8.6 %) SP-only (signal peptide, no TM), 59 (6.4 %) globular. No β-barrel calls were returned; DeepTMHMM is α-class-trained and a β-barrel-positive call at this length scale would be a soft observation in any case.

**Figure 5.**
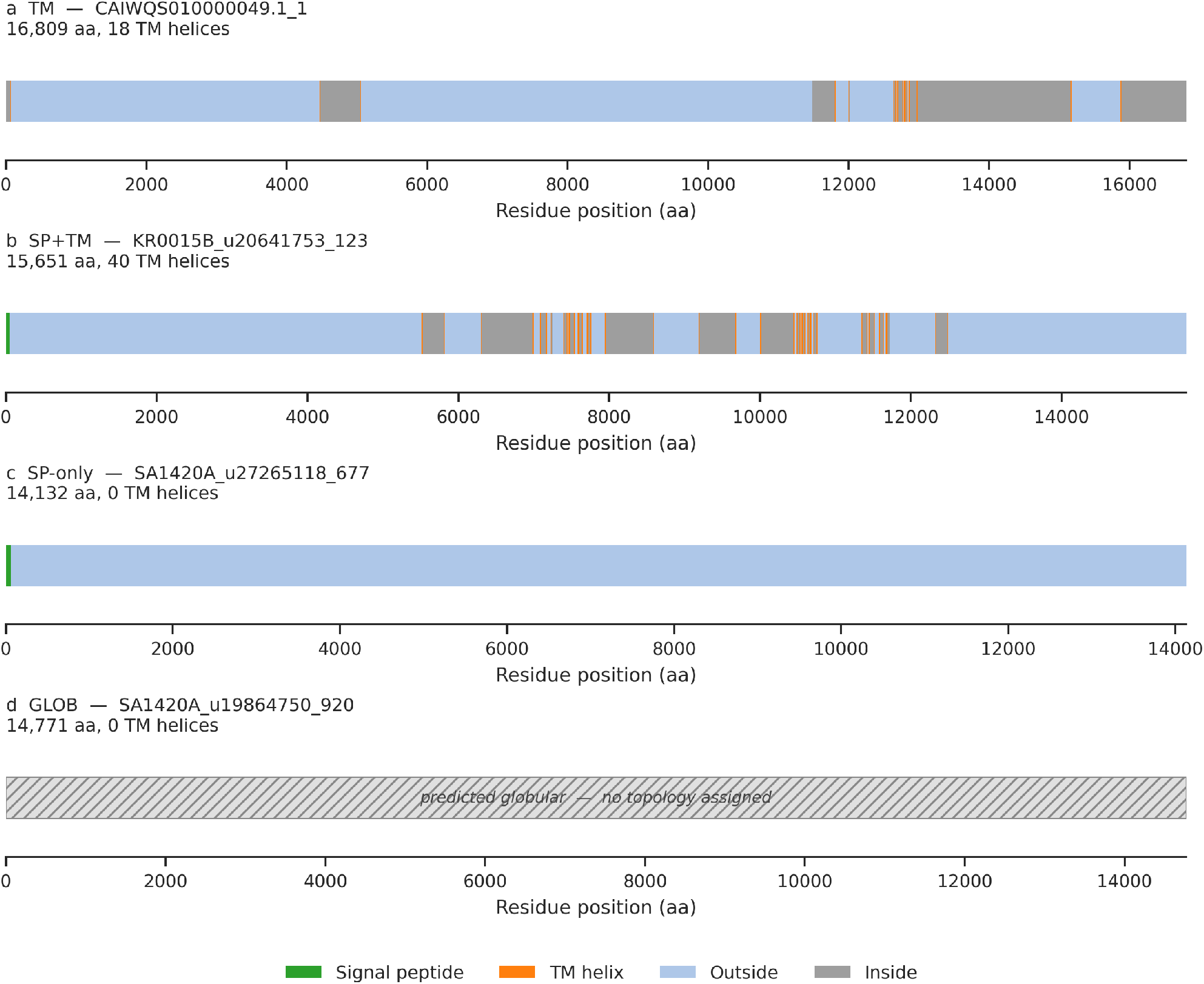
Membrane topology of the giant proteins. DeepTMHMM per-residue topology tracks for one representative protein per topology class: (a) TM (α-helical TM, no signal peptide; *n* = 422), (b) SP+TM (signal peptide plus TM helices; *n* = 356), (c) SP-only (signal peptide, no TM; *n* = 79), (d) GLOB (no TM, no signal peptide; *n* = 59). Representatives are the median-length protein in each class. Segments colored by feature type (signal peptide green, TM helix orange, outside light blue, inside gray); the GLOB panel is rendered as a single hatched bar to make clear that DeepTMHMM assigned no membrane topology — its default placement when no topology is predicted, not a positive call of an extracellular protein. All 916 long proteins screened; the four ≥ 100-kAA proteins were processed via the chunked-rescan and canonical-zone stitching pipeline described in Methods.

The membrane-bias intensifies sharply with length. At ≥ 30 kAA, 206 of 209 proteins (98.6 %) carry at least one TM helix or signal peptide, with three globular calls remaining. At ≥ 50 kAA (71 proteins) and across all 14 ≥ 80-kAA *Gorgyraeia* giants — 6 collection giants (4 above 100 kAA, 2 of 80–99 kAA) plus 8 giants on non-collection *Omnitrophota* contigs that the all-14-giant Pfam analysis already pulls in for §”Three architectural classes” — 100 % carry TM or signal-peptide content; the globular and SP-only fractions both drop to zero. Split across the 14: nine SP+TM, five TM-only, zero SP-only, zero globular. The four collection ≥ 100-kAA proteins are unanimous on this: three of four carry an SP+TM architecture (KR0015B_u4168876_114 at 147,155 AA with 147 TM helices, SA1420A_u33026402_205 at 132,636 AA with 157, SA1420A_u45987180_2000 at 104,139 AA with 68) and the fourth (SA1420A_u42255795_1875 at 122,534 AA) is TM-only with 331 TM helices. The 8 giants on non-collection *Omnitrophota* contigs split 5 SP+TM (KR0015B_u3039754_1735, KR0015B_u33178769_80, SA1420A_u23267945_853, SA1420A_u30763495_993, SA1420A_u6503474_406; 67–232 TM helices) and 3 TM-only (KR0015B_u10164256_188, SA1420A_u26689815_318, SA1420A_u42621477_744; 56, 62, 269 TM helices). TM-helix counts per protein scale with length — mean 22.2 helices at ≥ 10 kAA (median 16), 48.2 at ≥ 30 kAA, ~ 117 across the 14 ≥ 80-kAA giants — with Pearson r(length, TM count) = 0.61 across the catalog, a moderate correlation rather than a strictly linear scaling. The longest run of TM helices in a single polypeptide is 331 helices in `SA1420A_u42255795_1875` (122,534 AA), roughly one TM helix per 370 residues. Despite the absolute counts climbing, the per-residue *density* of TM segments stays essentially flat across length tiers at ~ 2 % of total residues (assuming canonical 20-AA helices). The giant proteins are therefore not denser membrane proteins than smaller ones; they are longer membrane proteins built to the same per-residue architectural recipe.

The half-and-half split between TM-only (46 %) and SP+TM (39 %) defines the dominant giant-protein architecture. TM-only is the canonical membrane-anchored-cytoplasmic-domain architecture — the bulk of the polypeptide predicted to sit on the cytoplasmic face of the membrane, threaded through it by the helices, with no signal peptide and no periplasmic targeting. At multi-kAA length the canonical Sec / SRP / YidC insertion mechanism that gives rise to this topology has not been experimentally validated, so the TM-only assignment here is a topology prediction rather than a confirmed architectural state. SP+TM is the canonical secreted-and-membrane-anchored architecture, with the N-terminal signal peptide directing the protein across the membrane and the TM helices then anchoring some portion of it back to the membrane on the periplasmic / outer face. The 9 % SP-only fraction is fully secreted across the inner membrane and not retained in it. These three classes together describe how 94 % of *Omnitrophota* long proteins reach the cell surface or the inner-membrane interface in some form. Pure cytoplasmic soluble enzymes are 6 % of the catalog, not the dominant reading. Restricting to the 14 ≥ 80-kAA giants sharpens the membrane-anchored bias further: all 14 *Gorgyraeia* giants — including the 147,155-AA protein that the Pfam analysis read as a multi-enzyme metabolic cluster — fall into the membrane-anchored classes, with no globular or SP-only call across the set.

To check that this collection-restricted topology pattern is not an artifact of CheckM2 curation, we ran DeepTMHMM on all 947 *Omnitrophota*-classified ≥ 10-kAA proteins in the 18-sample corpus — 427 on collection-genome contigs (205 contigs of the 229) and 520 on 308 non-collection *Omnitrophota* contigs that did not meet the collection’s CheckM2 bar but still bear long proteins (`scripts/ 56_chunk_corpus_omni_deeptmhmm.py`; proteins above the per-target length ceiling chunked and stitched identically). The headline survives the broader scale: 869 of 947 proteins (91.8 %) are membrane-targeted, with class breakdown 399 TM (42.1 %), 345 SP+TM (36.4 %), 123 SP-only (13.0 %), 78 globular (8.2 %), and 2 BETA. The two BETA calls — `KR0015B_u21703451_821` (15,129 AA) and `SA1420A_u2470583_1356` (10,556 AA) — are not canonical outer-membrane β-barrels but very long signal-peptide-bearing proteins on which DeepTMHMM places only short (5-residue) β-strand segments scattered between long periplasmic regions; the most plausible reading is a model artifact from forcing a 10–15-kAA secreted protein into one of the five DeepTMHMM topology classes, not the discovery of giant outer-membrane β-barrels. The length-tier intensification of the membrane bias is essentially the same at corpus scale as at collection scale (≥ 30 kAA 98.0 %, ≥ 50 kAA 100 %, ≥ 80 kAA 100 %); above 50 kAA the soluble fraction effectively disappears regardless of contig provenance. Within the corpus, the Baltic B10 *Omnitrophota* population (73 proteins) carries a substantially larger SP-only fraction (23 / 73, 32 %) and globular fraction (16 / 73, 22 %) than the Fennoscandian samples (~ 10 % SP-only, ~ 7 % globular). With *n* = 73 this is suggestive rather than established, but it is consistent with a Baltic water-column *Omnitrophota* population enriched in fully-secreted and soluble architectures relative to the deep-groundwater population.

### Genomic context of giant-protein-bearing loci

For each of the 14 ≥ 80-kAA giants from §”A multi-order capacity” — 6 collection circular contigs + 8 non-collection *Gorgyraeia* contigs — we examined the ± 20 ORF flanking-gene window and annotated those neighbors via KofamScan and mmseqs2 against UniRef90. The 6 collection flanks were pulled in the per-rotated-complete-genome Pyrodigal frame (`scripts/ 40b_giant_flanks_cohort_frame.py`), matching the frame the rest of this paper uses for collection gene IDs; the 8 non-collection flanks were pulled in the per-sample Pyrodigal frame (`scripts/ 40_giant_flanks_v2.py`) since their parent contigs are not rotated complete genomes. The pattern across both subsets is the same and recapitulates the West-Roberts et al. (2023) observation directly.

Across all 14 ± 20-ORF flanking windows (560 flanking ORFs total — 240 in per-rotated frame across the 6 collection giants, 320 in per-sample frame across the 8 non-collection giants; 79 % annotated):

- 42 flanking genes encode Type II / Type IV secretion-system or pilus-assembly components — 13 of the 14 giants carry at least one secretion / pilus flank. `SA1420A_u29588521_861` is a Type II secretion system GspF-domain protein at 52 % identity to *Candidatus* Ghiorseimicrobium GspF and a paralogous GspF (`SA1420A_u29588521_860`) hits *V. archaeovorus* Flp-family Type IVb pilin at 53 %; `SA1420A_u47349704_1588` is a Type II secretion system protein F at 42 % identity to *V. archaeovorus* T2SS-F; `SA1420A_u6503474_398` and eight surrounding flanks in the same locus collectively encode a Tad-pilus assembly cassette.
- 13 flanking genes encode glycoside hydrolases, with Glycoside Hydrolase family 2 catalytic-domain proteins (β-galactosidase / β-glucuronidase / β-mannosidase, the canonical extracellular polysaccharide-degrading family) dominant. The 147,155-AA protein’s flanking window itself carries two GH2 enzymes: `KR0015B_u4168876_128` at 59 % identity to *V. archaeovorus* β-galactosidase (UniRef90 `A0A410P2D0`) and `KR0015B_u4168876_130` at 74 % identity to *Candidatus* Sherwoodlollariibacterium GH2 (UniRef90 `A0A2G9YK91`). A non-collection flank `KR0015B_u10164256_195` hits the same UniRef90 Sherwoodlollariibacterium GH2 reference at 74 % identity, indicating the catalytic family recurs next to giants across collection and non-collection genomes alike.
- 24 flanking genes encode transposases or other mobile-DNA components, including IS110-family transposases (e.g. `SA1420A_u42255795_1888` at 93 % identity to a *Candidatus* Ratteibacteria IS110), IS5- and IS1634-family elements, integrases, and tyrosine recombinases. Several giants carry mobile-DNA-rich flanking neighborhoods, consistent with the broader MGE-integration pattern documented elsewhere in this paper.
- 6 flanking genes encode defense / restriction-modification machinery, all 6 on non-collection contigs (collection 0/240, non-collection 6/320) — including a MvaI/BcnI-family restriction endonuclease (`SA1420A_u42621477_763`), a ribbon-helix-helix antitoxin of a toxin–antitoxin system (`KR0015B_u3039754_1721`), and four flanking methyltransferases dispersed across three non-collection contigs.
- 2 flanking genes carry TonB-domain content, both on non-collection contigs (`KR0015B_u10164256_168` at 51 % identity to *Candidatus* Sherwoodlollariibacterium unditelluris TonB C-terminal domain; `SA1420A_u30763495_976` at 41 % to the same UniRef90 reference). Neither matches the canonical *Bacteroidetes* TBDR / *sus*-cluster polysaccharide-import architecture, which is therefore absent from both subsets. The giants are flanked by secretion / pilus and direct extracellular CAZyme machinery rather than by polysaccharide-import gene clusters.

The 6 collection giants and 8 non-collection giants show flank-category distributions that are similar on the secretion / pilus and CAZyme signals (T2SS/T4P flanks: 17/240 = 7.1 % in the collection set vs 25/320 = 7.8 % in the non-collection set; GH flanks: 7/240 = 2.9 % vs 6/320 = 1.9 %) and diverge on the mobile-DNA and defense signals (transposase flanks: 15/240 = 6.3 % vs 9/320 = 2.8 %; RM/ defense flanks: 0/240 vs 6/320 = 1.9 %). The shared T2SS / CAZyme axis is the dominant flank phenotype on both subsets; the collection vs non-collection divergence sits on the mobile-DNA / defense axis. The 8 non-collection contigs therefore largely represent the same secretion / CAZyme biology as the 6 collection giants and were excluded from the collection by CheckM2 quality heuristics, not by anything intrinsic to their giant-protein-bearing loci.

The flanking-gene complement is therefore consistent with the predator / cell-wall-degradation hypothesis (West-Roberts et al. 2023): 13 of 14 giants are flanked by secretion or pilus machinery, and the most recurring catalytic flanks are extracellular glycoside hydrolases (Fig. 6). Two flank features differ from prior reports: (i) the prominence of mobile-element flanks at a substantial fraction of giant-bearing loci, suggesting acquisition or movement of giant-protein loci on mobile cargo, and (ii) the carbohydrate-import signal being weaker than the secretion / CAZyme signal.

**Figure 6.**
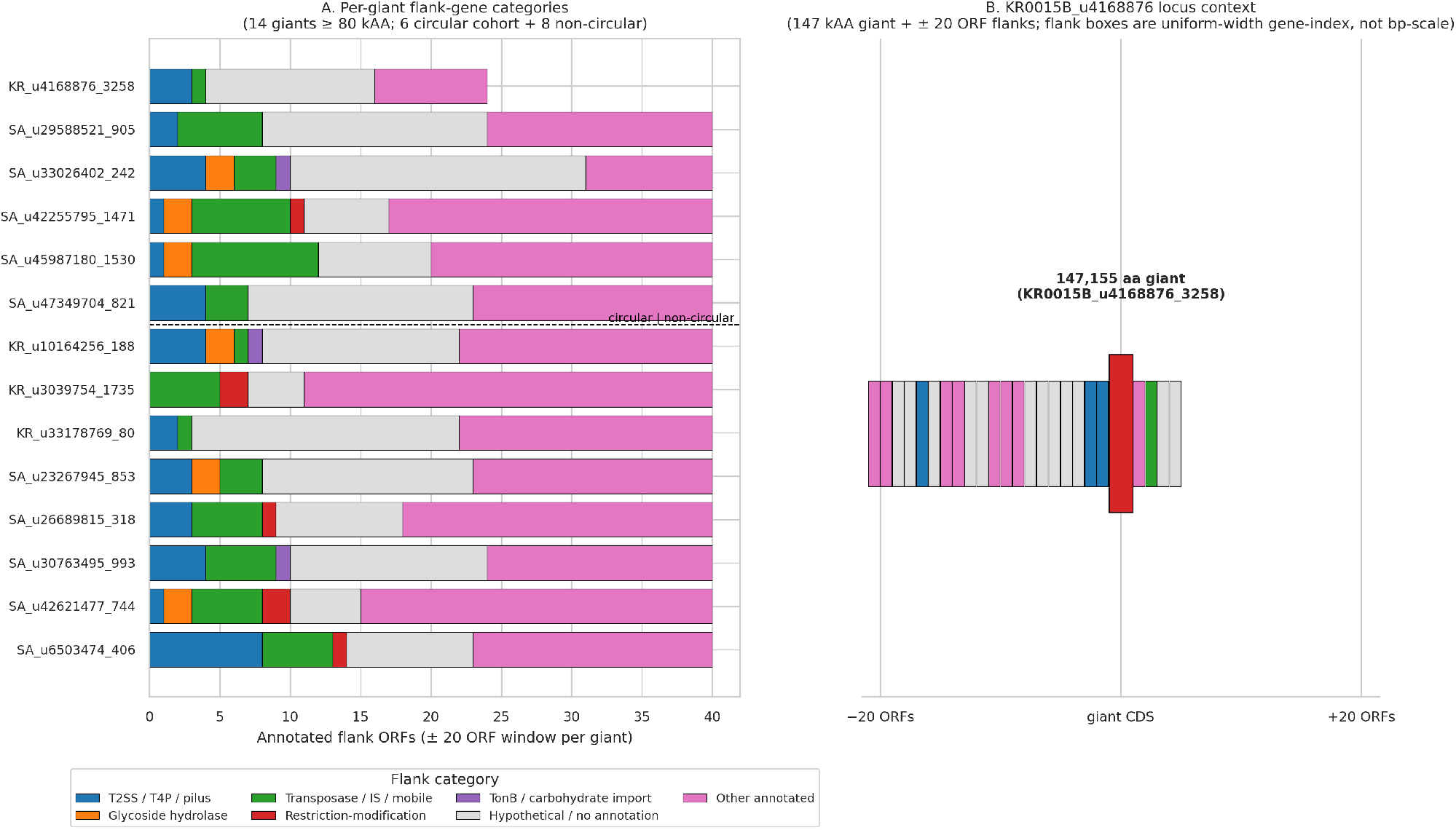
Genomic context. Neighborhood plots for the top giant-protein-bearing loci, highlighting Type II / IV secretion-system markers, glycoside hydrolase flanks, and transposase-rich neighborhoods. Canonical *Bacteroidetes* TonB-dependent receptor (TBDR) / *sus*-cluster polysaccharide-import architecture is absent from the collection (see §”Genomic context”); TonB-domain content is restricted to the 2 non-collection-contig flanking windows highlighted here.

### A constrained, host-dependent metabolic signature

The 176 complete-tier genomes show a constrained metabolic profile consistent with the host-dependent / episymbiotic lifestyle reported for the cultured *Gorgyraeia* representative *V. archaeovorus*. KofamScan annotations were cross-checked against the canonical KEGG KO list (KofamScan `ko_list`, downloaded 2026-02-27 from https://www.genome.jp/ftp/db/kofam/; 28,165 KOs); the pathway panel includes 103 KOs across central carbon metabolism, amino-acid biosynthesis, cofactor biosynthesis, muropeptide and cell-envelope synthesis, the respiratory chain, ATP synthase, and pathways previously flagged as informative for the lineage (West-Roberts et al. 2023).

### Central carbon — fermentative pattern, broken TCA

The lower half of canonical glycolysis is present in every genome — GAPDH (K00134), PGK (K00927), and enolase (K01689) all hit at 176/176 strict — and pfkA reaches 176/176 under relaxed threshold with UniRef90 confirmation. Hexose entry steps are partial across the collection: hexokinase 25/176, glucose-6-phosphate isomerase 67/176, with the rest carrying neither. The TCA cycle is fragmentary across all 176: IDH (K00031) and MDH (K00024) are PRESENT under relaxed thresholds with UniRef90 confirmation (165/176 and 166/176 respectively), citrate synthase (K01647) 21/176 relaxed, succinyl-CoA synthetase 5/176, fumarate hydratase 4/176, succinate dehydrogenase sdhA (K00239) 2/176 strict and sdhB (K00240) 2/176 strict, and 2-oxoglutarate dehydrogenase entirely ABSENT (K00164, K00658). The glyoxylate shunt is ABSENT in all 176 (K01637 ICL and K01638 aceB both 0/176). The signature is consistent with a fermentative central carbon metabolism that runs lower glycolysis to acetyl-CoA but does not close the TCA cycle and lacks anaplerotic glyoxylate input.

### Respiration — incomplete electron transport chain, absent aerobic terminal oxidase

Complex I (NADH-quinone oxidoreductase, *nuo*) is fragmentary across the collection: most subunits hit 10–43 / 176 genomes at strict thresholds — nuoF (K00335) and nuoE (K00334) reach 43 and 24 strict respectively; the remaining strand-A subunits cluster at ~11 strict; nuoG (K00336) is entirely absent — with no genome carrying the full operon. Complex II (succinate dehydrogenase / fumarate reductase) is essentially absent (sdhA K00239 at 2/176 strict, sdhB K00240 at 2/176 strict, sdhC K00241 at 0/176). The aerobic terminal cytochrome c oxidase aa3 (coxA / coxB / coxC) is ABSENT entirely (0/176 for all three subunits). F1F0 ATP synthase alpha and beta subunits are present at 125/176, but the gamma subunit reaches only 30/176 strict (35/176 relaxed), leaving most genomes without a complete F1F0 complex. There is no rhodopsin (K04641 bop and K04643 sop both ABSENT). The collection is therefore not configured for phototrophy, aerobic respiration through an aa3 terminal oxidase, or a complete F1F0 chemiosmotic apparatus.

### Amino-acid biosynthesis — selective

Tryptophan, histidine, and arginine biosynthesis are full pathways: trpE/G/D/A/B (K01657/K01658/K00766/K01695/K01696) all present at 174–176/176 strict; trpC (K01609) and trpF (K01817) cross strict thresholds in fewer genomes (143/176 and 164/176 strict respectively) but are PRESENT under relaxed thresholds (176/176 and 172/176) with UniRef90 confirmation; hisG/D/F/H/A/B/C/D similarly complete; argB/C/D, ornithine carbamoyltransferase (K00611), argG, and argH all 176/176 strict. Serine biosynthesis is partial — serA (K00058) 174/176 strict, but serC (K00831) only 9/176 and serB (K01079) 5/176 strict (26/176 relaxed), leaving most collection genomes without the canonical phosphoserine-aminotransferase / phosphatase route. The pattern matches a substrate that can produce aromatic and basic amino acids autonomously while drawing serine (and likely other simple amino acids) from environmental or host sources.

### Cofactor biosynthesis — partial

Thiamine biosynthesis: thiE / thiC / thiL / iscS all PRESENT (≥ 167/176); thiG (thiazole synthase, K03149) and thiD (HMP kinase, K00941) only 38/176 and 3/176 respectively. The biotin pathway is uniformly partial across the collection: bioF (K00652) 79/176 strict, bioA (K00833) 66/176, bioD (K01935) 74/176, bioB (K01012) 98/176 strict (106/176 relaxed) no genome carries the full set, and the pathway is a candidate marker for sub-clade differentiation. The selenocysteine machinery is incomplete — selB is PRESENT (174/176 relaxed) but selA (1/176) and selD (1/176) are essentially absent, suggesting that most genomes encode the elongation factor without the seryl-tRNA selenium transferase or the selenide:water dikinase needed to charge it.

### Cell envelope — canonical bacterial peptidoglycan biosynthesis is intact across the collection

The full cytoplasmic muropeptide pathway is present across the collection: glmS (K00820) 174/176; glmU (K04042), murA (K00790), murB (K00075), and murC (K01924) all 176/176 strict. D-alanine-D-alanine precursor supply — alanine racemase (K01775) and D-Ala-D-Ala ligase (K01921) — is also 176/176 PRESENT. Despite the ultra-small cells and the fragmentary energy metabolism, these genomes carry the complete classical bacterial peptidoglycan-biosynthesis machinery — not the wall-less L-form state sometimes adopted by small-genome host-dependent bacteria, and not the divergent peptidoglycan-light envelope chemistry historically described for other PVC lineages such as *Planctomycetota* and *Chlamydiota* (Jeske et al. 2015). They have conventional bacterial cell walls.

The metabolic profile across the 176 complete *Omnitrophota* genomes therefore describes a cell-walled bacterium with a fermentative central carbon metabolism, incomplete electron transport, no aerobic terminal oxidase, partial cofactor biosynthesis, and selective amino-acid biosynthesis. The combination full peptidoglycan machinery alongside extensively reduced energy and cofactor pathways — fits the host-dependent / predatory framing reported for the cultured *Gorgyraeia* representative *Velamenicoccus archaeovorus* (Kizina et al. 2022), where the bacterium attaches to and lyses methanogenic archaea and presumably draws cofactors, simple amino acids, and reduced carbon from the lysed host.

### Per-order resolution within the collection

(`scripts/50_metabolic_per_order.py`; complete-tier breakdown across the six orders with ≥ 12 genomes — *Gygaellales* 64, 2-01-FULL-45-10 16, *Tantalellales* 14, *Gorgyraeales* 14, *Velamenicoccales* 12, *Duberdicusellales* 12 — plus the 44-genome “other” bucket). Pathway-completeness scored as the number of panel KOs present at strict threshold in ≥ 50 % of order genomes is largely uniform: lower glycolysis 5/8, tryptophan / histidine / arginine biosynthesis at full completeness, muropeptide cytoplasmic synthesis 8/8, D-Ala-D-Ala precursors 2/2 in every order. The collection-wide deficits — fragmentary TCA (1/12 at strict ≥ 50 %), absent aerobic terminal oxidase (0/3), absent rhodopsin (0/2), partial Complex I (0/14 at strict ≥ 50 %, 4–7 of 14 at relaxed) — recur in every individual order. Two order-level deviations are visible at strict threshold but largely resolve at relaxed and so most likely reflect sequence divergence rather than pathway loss: *Gorgyraeales* F1F0 ATP synthase alpha and beta drop to 6/14 strict (vs 125/176 collection-wide) but recover to 14/14 relaxed, and *Gygaellales* biotin biosynthesis drops to 4/64 strict (vs 79/176 collection-wide) but reaches 61–64/64 relaxed across bioF / bioA / bioD / bioB. A separate comparison of *Pluralincolimonadales* (9 complete-tier genomes) against the rest of class *Gorgyraeia* (162 complete-tier genomes; `scripts/23_pluralincolimonadales_ko_profile.py`) recovers genuine metabolic differences rather than sequence-divergence patterns: *Pluralincolimonadales* loses 5’-methylthioadenosine phosphorylase, ribulose-phosphate 3-epimerase, acetate kinase, and V/A-type H+ / Na+-translocating ATPase subunits (present in 70–99 % of the rest of *Gorgyraeia* and ≤ 25 % of *Pluralincolimonadales*; all q ≤ 0.05), and gains glycerol-3-phosphate dehydrogenase subunit C (K00113; 8/9 vs 0/162). The collection-average profile reported above therefore describes the bulk of *Gorgyraeia* well, but smaller orders such as *Pluralincolimonadales* carry distinct metabolic specializations and should not be assumed to share the *V. archaeovorus* phenotype without per-order evidence. We do not interpret the partial pathways above as KofamScan false negatives: the cross-check confirms each call against the canonical KO definition and against UniRef90, and the high strict-hit rate in the pathways that ARE present (glycolysis lower half, Trp/His/Arg biosynthesis, muropeptide synthesis, cell-envelope precursors) indicates KofamScan’s profile thresholds are not systematically over-strict on this collection. Per-order metabolic-profile heatmap and full strict / relaxed / UniRef90 cross-check table are reported in Supplementary Figure S1.

### Hypervariable and mobile-cargo loci across 223 chromosomes

1,909 candidate hypervariable / mobile-element loci emerge from an assembly-level scan of the 229-genome collection — 223 of 229 chromosomes carry at least one (median 7 loci per genome, Q3 12, max 39; median span 11.5 kb, Q3 18 kb, max 109.5 kb; Fig. 7). The scan maps each genome’s own reads back to its assembly and identifies positions where soft-clip read termination piles up locally — the signature both of population-level cassette variability (where co-occurring strains diverge at a variable locus such as a polysaccharide cassette) and of integrative mobile elements (where reads support both an integrated and an episomal state, producing a “1.2× single-copy” interior coverage signal). The latter signature follows from the replicative-transposition mechanism, in which integrated and episomal copies coexist during a productive cycle (Harshey 2014); the chromosomal + episomal coverage arithmetic on the *u20424375* / *u17976357* worked example is in Nielsen (2026b). The peak-detection caller (`scripts/31_hvr_cohort_scan.py`) was calibrated against the *Omnitrophota* HVR previously documented on `KR0015B_u17976357` (Nielsen 2026b) — a 25-kb polysaccharide cassette at chromosome coordinates 547,000–572,000 bp, recovered by the caller as a 5.5-kb central peak at 559,000–564,500 bp with a peak ratio of 58× the genome-wide median.

**Figure 7.**
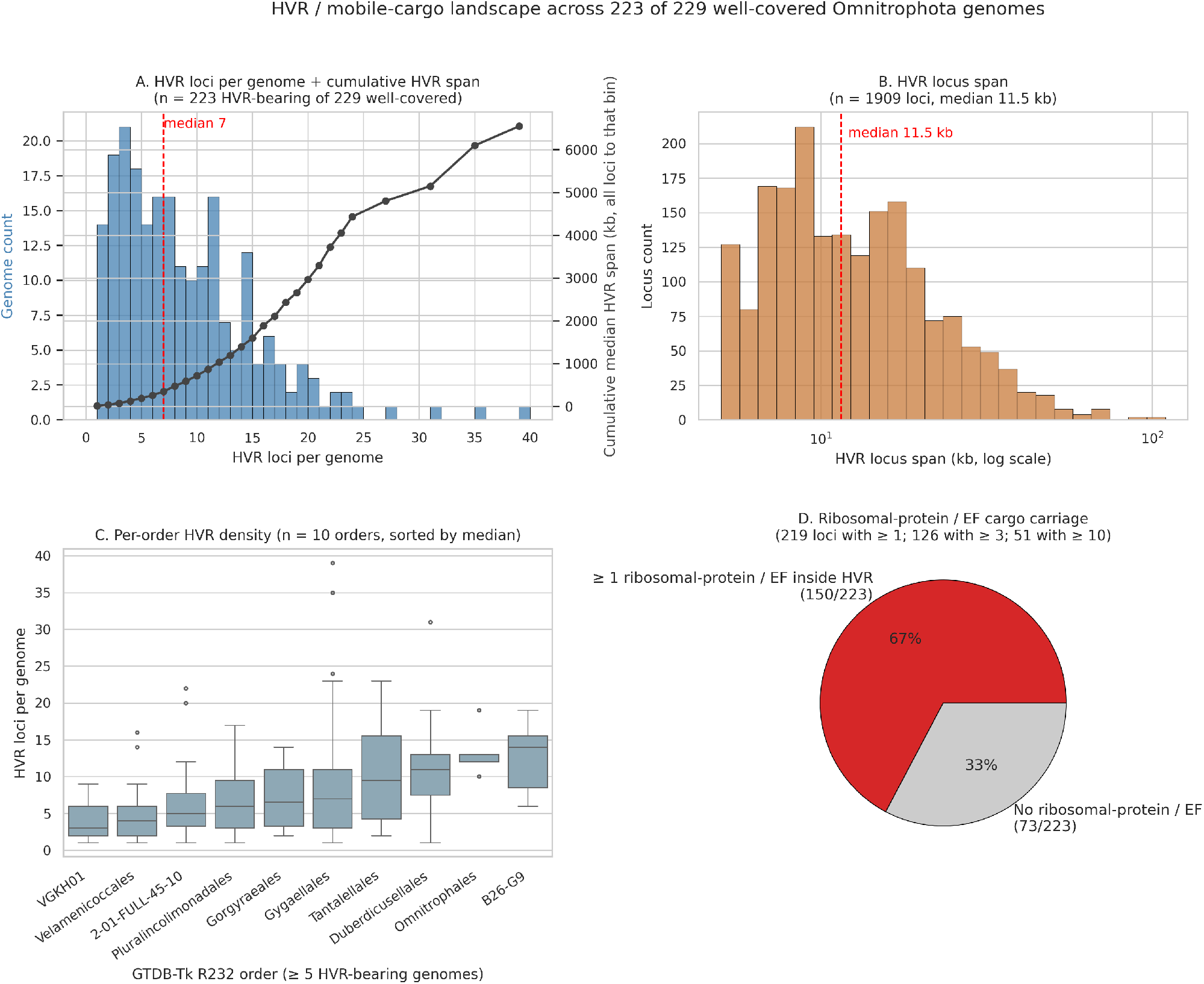
HVR / mobile-cargo landscape. Four-panel summary across the 223 of 229 genomes carrying at least one HVR call: (A) per-genome HVR locus count with cumulative HVR-span overlay, (B) per-locus span distribution on a log scale, (C) per-order HVR density (boxplots of HVR-loci-per-genome, GTDB-Tk R232 orders with ≥ 5 HVR-bearing genomes, sorted by median HVR-loci-per-genome), and (D) ribosomal-protein / EF-Tu / EF-G cargo carriage. The chromosome-level browser tracks for the *KR0015B_u17976357* + *u20424375* worked example are reported in Nielsen (2026b) and not duplicated here.

Functional annotation of all CDS overlapping these loci (`scripts/38_hvr_gene_content.py`; KofamScan and mmseqs2 against UniRef90 reach 44.9 % and 90.3 % annotation coverage respectively) recovers the canonical mobile-cargo gene complement. Site-specific DNA recombinases and IS-family transposases dominate (585 hits across four KO categories in the top-KO list), with type I and type III restriction-modification machinery (505 hits) and LPS / capsule glycosyltransferase, polysaccharide-biosynthesis, and gas-vesicle structural genes as the next-largest categories. The categorical breakdown (phage / mobile-element 4.3 %, restriction-modification and toxin-antitoxin 3.6 %, cell-surface biosynthesis 2.6 %, broad-substrate transport 1.8 %) maps cleanly onto the cross-replicon cargo catalog independently established for the same Fennoscandian deep-groundwater sample in Nielsen (2026b).

Three of those six categories (mobile-element movement, restriction-modification, cell-surface biosynthesis) are clearly prominent in the HVR cargo; two (toxin-antitoxin and T4P / secretion) are present but minor; one (metal / drug resistance) is essentially absent. The signal is therefore not a 1:1 reproduction of the Nielsen (2026b) catalog but the overlap is direct on the dominant categories.

The strongest signal is one the cross-replicon screen of Nielsen (2026b) deliberately could not see. Nielsen (2026b) filtered universal single-copy housekeeping markers — ribosomal proteins, EF-Tu / EF-G, RNA polymerase subunits, core replication and repair — out of the cross-replicon catalog to avoid deep-orthology false positives. All three clusters dropped from their 794-cluster cross-replicon cohort were ribosomal-protein / EF-Tu clusters carried on a single 233-kb integrative mobile element (`u20424375`) integrated into the `u17976357` chromosome across a 29-kb stretch at coordinates 635,000–664,000 bp — the only chromosomally-documented case in their dataset of an active MGE mobilizing most of a ribosomal-protein operon. Our HVR scan, operating on a different signal entirely (soft-clip peak detection at integration boundaries, not cross-replicon protein clustering), recovers the same housekeeping-marker class as prominent HVR content: 150 of 223 HVR-bearing genomes (67.3 %) carry at least one ribosomal-protein or EF-Tu / EF-G gene inside a called HVR, 126 of 1,909 HVR loci (6.6 %) host ≥ 3 ribosomal-protein / EF-Tu / EF-G cargo genes, and 51 loci host ≥ 10. Several chromosomes — including `KR0015B_u1208225`, `u10315151`, `u15895959`, `u17951743`, and `u11003791` — carry the canonical bacterial S10/spc operon (S12 → S7 → EF-G → EF-Tu → S10 → L3) in contiguous gene positions inside their HVR calls, in the same operon order Coenye and Vandamme (2005) report as the dominant bacterial S10/spc layout. Their single chromosomally-documented case is therefore not an isolated curiosity but the chromosomal manifestation of a recurring phenomenon: ribosomal-protein operons recur inside HVR windows at substantial frequency across the collection. Whether each individual ribosomal-protein operon-bearing HVR represents a fully integrated MGE — as in the *u20424375* / *u17976357* worked example — or a population-variable locus with operon-internal heterogeneity is not resolved by the HVR signal alone, and we return to it in the next paragraph.

HVR density varies roughly threefold across the orders represented in the 223 HVR-bearing genomes (complete and high-quality combined): *Velamenicoccales* (median 4 loci per genome, n = 19 of 19 in collection) and *Pluralincolimonadales* (median 6, n = 15 of 17) are HVR-poorest, while *Duberdicusellales* (median 11, n = 15 of 15), *Omnitrophales* (median 12, n = 5 of 5), and *B26-G9* (median 14, n = 8 of 8) are HVR-richest. Several smaller orders show median densities of 10–21 with sample sizes of 1–3 and are not yet well powered. *Velamenicoccales* — the order containing the cultured episymbiont *V. archaeovorus* — sits at the HVR-poor end of the distribution despite every genome in the order carrying at least one ≥ 10-kAA giant protein, suggesting giant-protein burden and HVR burden are not coupled at the order level.

The detection scheme cannot distinguish active integrative mobile elements — where a cassette is present in both integrated and episomal copies in the read pool — from population-variable surface cassettes — where co-occurring strains carry divergent variants of the same locus. For the previously-documented `u17976357` case (Nielsen 2026b) both phenomena co-localize on the same 2.1 Mbp chromosome: the 25-kb polysaccharide variable cassette at coordinates 547,000–572,000 bp and the 29-kb integrated mobile element at coordinates 635,000–664,000 bp are separate but neighboring features ~60 kb apart. The HVR caller fires preferentially on the polysaccharide cassette — heterogeneous reads producing a strong soft-clip peak — and not on the high-identity integrated mobile-element backbone, consistent with the 99.3 % integrated-versus-episomal identity reported by Nielsen (2026b). The 1,909 calls thus likely include both classes; teasing the two phenomena apart per locus is a methods-paper follow-up.

## Discussion

### Complete-genome collection

The 176 new complete genomes expand the prior GTDB R232 inventory of two complete *Omnitrophota* assemblies by 88-fold. This makes pangenome structure, replication-origin context, integration-site distribution, and population-level questions tractable at chromosome resolution across the phylum. All 176 were produced by a single assembler pass on Oxford Nanopore long reads (myloasm; Shaw et al. 2026) with no manual sequence editing — the workflow is fully reproducible.

### A giant-protein phenotype concentrated in class *Gorgyraeia*

*Omnitrophota* host many very long proteins, and the long end of the corpus-wide length distribution is class-restricted to *Gorgyraeia*. The 14 longest examples in the corpus — all above 80 kAA — are on 14 distinct *Gorgyraeia* contigs across multiple *Gorgyraeia* orders, pairwise divergent in sequence by all-vs-all DIAMOND. The one full-length cross-pair alignment across these 14 is between a 122-kAA and a 99-kAA *Gorgyraeaceae* giant at 58.6 % identity over essentially the full length, indicating that the giant-protein architecture can be conserved across divergent members of the same family. A single such pair does not on its own establish family-wide conservation; we report it as proof-of-principle. At the broader ≥ 10 kAA threshold the phenotype is dense and order-spanning: ~24–28 % of GTDB-Tk-classified *Omnitrophota* contigs in our Fennoscandian deep-groundwater and Baltic Sea samples host at least one ≥ 10-kAA protein. Both findings — class-level concentration at the longest end of the length distribution and enrichment across the 12 of 20 *Omnitrophota* orders with ≥ 3 representatives in the collection — establish the long-protein phenotype at collection scale on independently assembled material (West-Roberts et al. 2023). The collection is biased toward Fennoscandian deep groundwater (170 of 176 complete genomes from KR0015B + SA1420A); claims here describe *capacity* in the orders sampled, not phylum-wide *distribution*.

What these proteins do at the level of biochemistry is not fully resolved by the present data, but the DeepTMHMM topology survey of §”Membrane topology of the giant proteins” sharply narrows the architectural space. The flanking-gene complement — secretion / pilus machinery and extracellular glycoside hydrolases recurring next to giants — is most easily read as extracellular cell-wall-targeting effector function (West-Roberts et al. 2023). The DeepTMHMM data are consistent with that reading for a substantial fraction of the collection: 356 of 916 long proteins (39 %) carry both a signal peptide and TM helices (SP+TM architecture), the canonical secreted-and-membrane-anchored topology, and a further 79 (9 %) are SP-only and fully secreted. The dominant architecture is something different but still membrane-associated: 422 of 916 (46 %) are TM-only — no signal peptide, multiple TM segments — the canonical membrane-anchored-cytoplasmic-domain phenotype, where the bulk of the polypeptide remains on the cytoplasmic face of the inner membrane and the TM helices anchor it there. Pfam Module 1 (signal-transduction / regulatory architecture: methyltransferase, histidine-kinase / HSP90-like ATPase, GNAT acetyltransferase, HAD hydrolases) and Module 2 (sugar metabolism: starch synthase, glycosyl transferase group 1, phosphoglucomutase, response-regulator receiver, radical SAM) are entirely compatible with the TM-only inner-membrane-anchored reading, where the regulatory and metabolic content sits on the cytoplasmic face. Pure cytoplasmic soluble enzymes — globular proteins with neither a signal peptide nor a TM helix — are 6 % of the catalog (59 of 916), a minority architecture rather than the dominant one. The 147,155-AA protein that the Pfam analysis read as a multi-enzyme metabolic cluster is, by DeepTMHMM, an SP+TM protein with 147 TM helices — secreted across the inner membrane via its N-terminal signal peptide and anchored to it by the helical bundle, not an intracellular soluble cluster. The other three ≥ 100-kAA proteins are similarly membrane-associated, two SP+TM and one TM-only. The intracellular-metabolic-cluster reading we floated earlier for that protein is therefore contradicted by its topology; the multi-Pfam metabolic content sits on the periplasmic / extracellular face.

With that revision the surviving hypotheses are: (i) extracellular cell-wall-targeting effectors, supported for the SP+TM and SP-only fractions including the 147,155-AA protein; (ii) secreted multi-domain CAZymes operating on extracellular polysaccharide substrate, also supported for the SP-containing fractions; (iii) inner-membrane-anchored cytoplasmic-domain enzymes — supported for nearly half the catalog, the TM-only fraction; (iv) integrative-element passenger genes acquired and propagated by the mobile-cargo machinery documented in §”Hypervariable and mobile-cargo loci across 223 chromosomes”. These are not mutually exclusive, and our three repeat-architecture classes multi-enzyme metabolic, tandem-repeat scaffold, dockerin-bearing — likely distribute across them. What remains for follow-up is targeted heterologous expression for the tandem-repeat and dockerin classes and structural prediction for the dark-matter fraction.

### A constrained metabolic profile consistent with the cultured *Gorgyraeia* phenotype

The collection metabolic profile fits the host-dependent / episymbiotic lifestyle established for *V. archaeovorus*: fermentative central carbon, a broken TCA cycle, no aerobic terminal oxidase, an incomplete F1F0 chemiosmotic apparatus, partial cofactor biosynthesis with notable deficits in biotin and selenocysteine, and selective amino-acid biosynthesis with full Trp / His / Arg pathways alongside partial serine and absent glyoxylate-shunt input. These reductive features sit next to a fully intact bacterial peptidoglycan-biosynthesis machinery. The combination is consistent with a substrate that draws cofactors and simple carbon from a host while retaining its own canonical bacterial cell wall. The per-order resolution we report is consistent with that framing across the six largest orders (≥ 12 complete-tier genomes each): pathway-completeness is uniform across the pathways above, and the few order-level deviations at strict KofamScan thresholds (*Gorgyraeales* F1F0, *Gygaellales* biotin) recover at relaxed thresholds and so most likely reflect sequence divergence rather than pathway loss. Smaller orders behave less uniformly: *Pluralincolimonadales* (9 complete-tier genomes) has genuine metabolic specializations — losses of methylthioadenosine phosphorylase, ribulose-phosphate 3-epimerase, acetate kinase, and V/A-type ATPase subunits, plus a gain of glycerol-3-phosphate dehydrogenase subunit C that distinguish it cleanly from the bulk *Gorgyraeia* baseline. The collection-average profile is therefore informative for the dominant orders, but the smaller *Omnitrophota* orders should be treated as separate phenotypes pending order-by-order biochemical evidence. Two limitations bound the inference: the collection is Fennoscandian-deep-groundwater dominated, so the collection-level profile may not generalize to *Omnitrophota* in surface freshwater or host-associated habitats where the phylum is also reported but underrepresented in our samples; and KofamScan profiles describe potential — what enzymes the genomes can produce — not flux. Pairing the metabolic substrate with the giant-protein flanking analysis gives a coherent host-dependent / predatory cell-biology framing, but flux measurements remain a target for downstream work on the cultured *V. archaeovorus* representative or a future high-quality cultured isolate.

### Integration and variability loci across the collection, and a recurring ribosomal-operon-cargo signal

Mapping each complete chromosome’s own reads back to its assembly and calling positions where soft-clip read termination piles up locally — the long-read locus-level analogue of the short-read mobile-genetic-element (MGE) integration signal first systematized by Durrant et al. (2020) — flags 1,909 candidate loci across 223 of 229 chromosomes. We call these loci “HVRs” here as a generic label; biologically the 1,909 calls are heterogeneous, spanning polysaccharide-cassette variability of the *Pelagibacter* type, active mobile-element integration boundaries, restriction–modification (RM) islands, and capsule clusters in a single class. The detection signal is the same; the underlying biology is not.

The cargo composition recovered by per-locus KofamScan and mmseqs2 against UniRef90 maps cleanly onto the cross-replicon mobile-cargo catalog independently established for the same deep-groundwater samples (Nielsen 2026b): site-specific recombinases and IS-family transposases, type I and type III restriction–modification systems, and capsule and LPS glycosyltransferases. Canonical bacterial S10/spc ribosomal-protein operon content appears inside HVR windows on 150 of 223 HVR-bearing genomes (67 %). That housekeeping-cargo class is the one Nielsen (2026b) had to filter out of their cross-replicon catalog to avoid deep-orthology false positives, with a single chromosomally-documented case (`KR0015B_u17976357` hosting the integrated *u20424375* mobile element across a 29-kb stretch at coordinates 635,000–664,000 bp). Our scan, operating on a different signal entirely, recovers that case across the collection.

The 150/223 ribosomal-cargo signal could in principle be an artifact of cross-genome read mismapping in a highly-conserved cargo class — reads from near-relatives landing on the same chromosomal locus and producing a soft-clip pile-up without actual integration. The mapping design rules that mechanism out directly: every genome was mapped against its own reads only, not against a pooled or cross-sample read set, so cross-genome mismapping has no opportunity. Two further observations argue against any residual artifact: the `u17976357` worked example carries the ribosomal-protein operon inside a mobile-element backbone with characteristic boundary breakpoints whose positions are unrelated to the per-protein conservation ranking of the operon’s content; and the 51 loci with ≥ 10 ribosomal-protein / EF cargo genes carry the genes in canonical bacterial S10 / spc operon order (S12 → S7 → EF-G → EF-Tu → S10 → L3) — operon synteny is a positive indicator of integrated genuine biology rather than of conservation-similarity mismapping, which would not respect gene order.

We explicitly tested whether the *u20424375* / *u17976357* element of Nielsen (2026b) classifies as a Mu-class transposable bacteriophage — replicative-transposition phages with the canonical DDE_2 transposase plus a five-protein GemA / Mor / portal / head-tail-connector / virion-morphogenesis architecture — by the six conserved-marker definition of Zhang et al. (2023). It does not: only one of six markers is present, the IS240-family transposase, which Zhang accept only as a fallback for the DDE_2 marker in combination with the other five, and no GemA, Mor, portal, head-tail connector, or virion-morphogenesis protein is on either *u17976357* or *u20424375*. The two-state coverage signal is therefore consistent with general integrative-mobile-element behavior but is not mechanistically attributable to Mu-class replicative transposition. The collection puts a rate on the phenomenon: ribosomal-protein operons appear as HVR-window content on 150 of 223 HVR-bearing *Omnitrophota* chromosomes (67 %). The mechanism is not resolved — the element carries neither the integrase / conjugation machinery of classical ICEs nor the phage-structural markers of Mu-class transposable bacteriophages.

## Methods

### Samples and sequencing

Oxford Nanopore long-read metagenomes from 18 *Omnitrophota*-candidate samples: two Fennoscandian deep-groundwater sites and the 16-sample Baltic Sea water column. Deep-groundwater sites: KR0015B at 69 m below sea level and SA1420A at 201 m below sea level, both at the Äspö Hard Rock Laboratory, Sweden. Baltic samples: B1–B12, B14, B15, B17, B19. Raw Oxford Nanopore reads are deposited under NCBI BioProject `PRJNA1461169` (Fennoscandian groundwater; samples KR0015B and SA1420A) and `PRJNA1461195` (Baltic Sea water column; the 16 Baltic samples).

### Assembly

myloasm v0.3.0 with `-b 250` (non-default Bloom-filter size, chosen primarily to increase memory headroom on the deep-coverage Fennoscandian samples). Per-unitig circularity, length, and read coverage were read from the assembler’s `<sample>_info.tsv` output; coverage is taken as `Depth1` (reads aligning at ≥ 99 % identity). We use “contig” throughout the body of this paper as a synonym for myloasm’s “unitig” (a maximally unambiguous walk through the assembly graph); “unitig” is retained in Methods only where it labels the assembler’s per-unitig output columns and the corresponding script names.

### Quality and topology

CheckM2 v1.1.0 (Chklovski et al. 2023) with the bundled `uniref100.KO.1.dmnd` reference database (2025-02-08 build) for completeness and contamination. Circular topology from myloasm’s `*_info.tsv` per-unitig flag. **Complete** = single contig + CheckM2 ≥ 90 % completeness + < 5 % contamination + assembler-circular flag. **High quality** = single contig + CheckM2 ≥ 90 % / < 5 %, not assembler-circular. **Contig** = anything below the high-quality bar. In our pipeline the single-contig and not-circular qualifiers are implied by *high quality*; this paper never repeats them when the term has already been introduced.

### Rotation to origin

Complete genomes were rotated so that the *dnaA* (chromosomal replication initiator) start codon lands at position 1 of the forward strand. *dnaA* identification is anchored on per-genome MMseqs2 hits against UniRef90 (`clusters/all_vs_uniref90.tsv`) rather than KofamScan K02313, which is unreliable for divergent *Omnitrophota dnaA*; the highest-coverage gene whose UniRef90 hit description matches `dnaA` or `chromosomal replication initiator` is selected. The rotation chain is `scripts/04_rotate.py` (paper-local driver) → `rotate_v2.py` (project-shared engine). For minus-strand *dnaA*, the chromosome is reverse-complemented before the coordinate shift so that *dnaA* ends up on the forward strand. Genomes whose assembly seam interrupts the *dnaA* CDS — manifesting as a fragmented or partial *dnaA* ortholog at the contig boundary — are repaired in a separate pass (`scripts/09_repair_dnaA_one.py`, calling the project-shared `repair_dnaA.py`) before re-rotating. Rotation outcomes are verified by `scripts/05_verify_rotation.py`, which re-calls genes on the rotated assembly with Pyrodigal and confirms that gene_1 is a single-CDS *dnaA* starting at position 1 on the forward strand. After rotation, gene calls are re-done with Pyrodigal under the rotated regime (`force_nonsd=False`).

### Taxonomic placement

GTDB-Tk v2.4.0 (Chaumeil et al. 2022) `classify_wf` against GTDB release R232, using the bundled bac120 and ar53 marker placements, AAI screen, and pplacer reference-tree placement. For *Omnitrophota* genomes carrying multi-kAA proteins, the bundled HMMER `hmmalign` step refuses targets longer than 100,000 amino acids; the 147,155-AA protein and the other multi-kAA giants are not bac120 markers and the failure is silent for placement purposes, but we patched the local install to skip oversize targets at the alignment step rather than abort the per-genome run.

### Gene prediction

Pyrodigal v3.6.3 (Larralde 2022; underlying Prodigal algorithm: Hyatt et al. 2010) was run in two regimes. (i) Per-genome single-mode (`meta=False`, train on the concatenated genome, then `find_genes` per contig) for the 229 collection genomes plus 8 reference proteomes (4 NCBI *Omnitrophota* + 4 sample-derived PVC outgroups). (ii) Per-sample metagenomic mode (`meta=True`) for the cross-sample long-protein scan across the 18 *Omnitrophota*-candidate samples; in meta mode there is no per-genome SD training step, so the `force_nonsd` flag is not applicable and is not set in the driver. In the per-genome regime, the `force_nonsd` flag was set per-genome to match assembly state — `force_nonsd=False` for rotated complete genomes (where the assembly carries natural upstream context after rotation to ORC1/dnaA at position 1), and `force_nonsd=True` for unrotated reference genomes (the project default; avoids over-extension of gene_1 when chromosomal upstream context is absent).

Long ORFs are not filtered by default: Prodigal caps the minimum predicted ORF length at 90 bp but imposes no maximum, so the 147,155-AA protein and the other multi-kAA giants discussed in Results are direct Pyrodigal output.

### Phylogenomics

OrthoFinder v3.1.4 (Emms & Kelly 2019) in default mode (DIAMOND all-vs-all + MCL clustering + dendroblast-based species-tree-aware orthology) was used for single-copy orthogroup identification on 184 input proteomes (176 complete-tier genomes + 4 NCBI *Omnitrophota* reference proteomes including *V. archaeovorus* GCF_004102945.1 + 4 sample-derived PVC outgroup proteomes). MSA-based and IQ-TREE-based gene-tree modes (`-M msa`, `-T iqtree3`) were not used. The 97 strict single-copy orthogroups were re-aligned with MAFFT L-INS-i (`--localpair --maxiterate 1000`), trimmed with BMGE v2.0 (`-t AA -m BLOSUM30 -h 0.5 -g 0.5 -b 3`), concatenated, and inferred under IQ-TREE v3.0.1 (`-T 32 -m LG+F+R10 -B 1000 -alrt 1000 -bnni --seed 42 -o 1W_u26197113,1W_u2824098,8W_u1250509,B7_u100762`). For the extended phylogenetic placement against all GTDB R232 *Omnitrophota* species representatives (Supplementary Figure S2), we used the cohort-HMM recruitment pipeline of Nielsen (2026a). The 97 cohort-OG alignments were converted to profile HMMs; `hmmsearch` was used to recruit the corresponding ortholog from each NCBI proteome; per-OG gene-tree QC classified each recruited tip relative to the cohort MRCA descendant set on a per-OG basis; and a per-MAG paralog-rate filter excluded MAGs above a 25 % per-MAG paralog-candidate rate before supermatrix concatenation. The recruitment step is necessary because the OrthoFinder + DIAMOND baseline drops 175 of 710 evaluable NCBI MAGs at a ≥ 90 % gap cutoff against the 97-OG cohort supermatrix despite CheckM2 considering all 175 HQ (median completeness 94 %), with contig count and missing-marker count essentially uncorrelated across the 710 candidates (Spearman ρ = 0.03). The drop reflects the cohort-fit character of our 97-OG marker set and the depth of DIAMOND-similarity attrition at phylum scope, not GTDB-level quality; HMM profile detection supplies the recruitment power that pairwise DIAMOND cannot reach, and the per-OG gene-tree QC supplies the orthology call on a signal independent of the recruitment. The resulting 938-tip supermatrix (229 in-house cohort + 4 PVC outgroups + 705 GTDB R232 species representatives) was inferred under identical IQ-TREE parameters as the 184-tip tree. Headline phylogenetic claims rest on the 184-tip collection tree; the 938-tip tree is taxonomic placement only. Class-level names in this paper follow GTDB R232 (*Gorgyraeia, Omnitrophia*); R226 used the placeholder `Koll11` for what R232 renamed to *Gorgyraeia*, so any per-sample GTDB-Tk output predating R232 retains the placeholder in the raw data and is translated to the R232 name in all body claims.

### KEGG ortholog annotation

KofamScan v1.3.0 (Aramaki et al. 2020) `exec_annotation`, run per-genome on the Pyrodigal protein sets via `scripts/12b_kofamscan_per_genome.py`. The KEGG ortholog profile distribution and `ko_list` threshold file were downloaded 2026-02-27 from https://www.genome.jp/ftp/db/kofam/. Both strict (above the per-profile adaptive threshold) and the relaxed e-value class were retained at the script level; only strict hits are reported in the body of this paper. KO-to-name resolution is loaded from the canonical `ko_list` at runtime rather than from any source-embedded dictionary, and every annotation referenced in the metabolic discussion is re-resolved against `ko_list` via `scripts/39_kofamscan_crosscheck_complete.py` (the paper-local adapter; the canonical engine is the project-shared `kofamscan_crosscheck_v2.py`) before reporting.

### Long-protein scan

`scripts/15_long_proteins_cross_sample.py` walks the 18 sample-level Pyrodigal metagenomic calls (`meta=True`; `force_nonsd` not set as it is meta-mode-vacuous) across the *Omnitrophota*-candidate sample panel and emits every protein at or above 10,000 amino acids to `logs/ long_proteins_cross_sample_ge10kaa.tsv` with sample, unitig, gene_id, length, and start/end coordinates. Length tiers used in the body — ≥ 10 kAA, ≥ 30 kAA, ≥ 80 kAA — are downstream slices on this table, not separate scans. Per-protein taxonomy is joined against the per-sample GTDB-Tk R226 contig classifications via `scripts/16_long_proteins_taxonomy.py`; the per-sample runs predate R232 and carry the R226 placeholder class name `Koll11` for what R232 renamed to *Gorgyraeia*. Class-level claims in this paper use the R232 name throughout. *Omnitrophota*-contig denominators (used for the “X % of *Omnitrophota* contigs host a ≥ 10-kAA protein” rates) come from `scripts/17_omni_unitig_counts.py`, which counts classified-*Omnitrophota* contigs per sample from the same GTDB-Tk output.

### Domain architecture

InterProScan v5.59-91.0 (Jones et al. 2014) restricted to an architectural member-database panel of Pfam, SMART, CDD, PANTHER, ProSitePatterns, ProSiteProfiles, Gene3D, Coils, and MobiDBLite was run on the 912 proteins at ≥ 10 kAA (`scripts/24_interproscan_long_proteins.py`). SUPERFAMILY was dropped from the panel: the bundled hmmlib_1.75 (October 2022 release) carries duplicate model accessions that hmmpress 3.4 rejects with “secondary keys not unique”; not fixable without rebuilding the HMM library upstream. Pfam + Gene3D + SMART + CDD between them cover the architectural classification SUPERFAMILY would have contributed. Signal-peptide and TM-helix prediction is handled separately by DeepTMHMM (below). The InterProScan precalculated-match lookup service was disabled (`-dp`) so that every input is annotated from local databases. Domain co-occurrence and per-protein architecture summaries were generated by `scripts/ 26_ips_domain_cooccurrence.py` and `scripts/47_fig4_domain_architecture.py`. InterProScan’s bundled HMMER scan refuses individual targets longer than 100,000 amino acids, so the 13 proteins above that cap were split into ≤ 100-kAA non-overlapping chunks, re-scanned separately, and re-projected back to the parent protein’s coordinates for the architecture summary. Transmembrane-helix prediction used DeepTMHMM v1.0 (Hallgren et al. 2022) in the standard Academic local-install configuration. The 912 ≤ 99,999-AA long proteins were split into four length-balanced batches (228 sequences each, greedy-bin-packed by descending length so total residue load per batch is matched to within ~ 0.01 %) and run in parallel on CPU with 16 threads each (`deeptmhmm/split_batches.py`, sibling of `scripts/`). Total wallclock was ~ 12 h per batch (~ 6 h ESM-2 embedding + ~ 5 h topology decoding), running concurrently. DeepTMHMM applies an internal per-target length limit empirically bounded above ~ 85 kAA in the ESM-1b / fair-esm 0.4.0 path bundled with the Academic install; the four collection ≥ 100-kAA giants were therefore chunked into ≤ 80,000-AA pieces with 5,000-AA overlap (`scripts/54_chunk_giants_deeptmhmm.py`), predicted per-chunk under the same DeepTMHMM configuration, and stitched back into full-length per-residue tracks using a canonical-zone rule that attributes each parent residue to exactly one chunk (half-overlap exclusion at interior boundaries, full extent at protein edges; `scripts/ 55_stitch_giants_deeptmhmm.py`). Signal-peptide (‘S’) calls on non-first chunks are downgraded to ‘O’ before stitching, since a signal peptide can only exist at the protein’s N-terminus and any ‘S’ on a later chunk is a chunking artifact. After stitching, protein-level topology classes are re-derived from the unified track using DeepTMHMM’s own rule (presence of ‘S’ and ‘M’ → SP+TM, ‘S’ alone → SP, ‘M’ alone → TM, ‘B’ alone → BETA, neither → GLOB). All 916 proteins in the domain-architecture catalog are covered. The 8 ≥ 80-kAA giants on non-collection *Omnitrophota* contigs that the all-14-giant Pfam analysis already pulls in for §“Three architectural classes” — `KR0015B_u10164256_188` (81 kAA), `KR0015B_u3039754_1735` (94 kAA), `KR0015B_u33178769_80` (93 kAA), `SA1420A_u23267945_853` (85 kAA), `SA1420A_u30763495_993` (90 kAA), `SA1420A_u42621477_744` (99 kAA), `SA1420A_u6503474_406` (80 kAA), and `SA1420A_u26689815_318` (115 kAA) — were processed identically via `scripts/ 54b_chunk_open_giants_deeptmhmm.py` so that the 14-giant topology analysis runs end-to-end on a single consistent pipeline.

For corpus-scale topology coverage extending beyond the collection, DeepTMHMM was additionally run on all 947 ≥ 10-kAA proteins on GTDB-Tk-classified *Omnitrophota* contigs in the 18-sample corpus (`scripts/56_chunk_corpus_omni_deeptmhmm.py`), with 14 proteins above the per-target ceiling chunked and stitched as above. The corpus-scale set is derived from the per-sample metagenomic Pyrodigal calls (the same input frame as `scripts/ 15_long_proteins_cross_sample.py`) and extends topology coverage to 308 non-collection *Omnitrophota* contigs that the collection run does not address. Total wallclock for the corpus-scale run was ~ 16 h on 4 parallel 16-thread workers.

### HVR / mobile-element scan

Each complete genome’s own Oxford Nanopore reads were mapped back to its rotated assembly with minimap2 (`-ax map-ont`). Soft-clip read termini were counted in non-overlapping windows. Peaks were called where the per-window soft-clip rate exceeded both an absolute clip count and a multiplier over the local flanking median. Caller thresholds (`scripts/31_hvr_cohort_scan.py`) were calibrated against the *Omnitrophota* HVR on `KR0015B_u17976357` (Nielsen 2026b): MIN_PEAK_TOTAL_CLIPS 30 per window, MIN_PEAK_RATIO 10× over flanking median, MIN_LOCUS_PEAK_RATIO 50× the genome-wide median at the locus level, MIN_LOCUS_TOTAL_CLIPS 500, MIN_LOCUS_KB 5.0, MIN_COV_DROP_VS_FLANK 1.2. Locus gene content was joined against KofamScan (KEGG `ko_list`, downloaded 2026-02-27 from https://www.genome.jp/ftp/db/kofam/) and mmseqs2 vs UniRef90 via `scripts/ 38_hvr_gene_content.py`. The signal is the long-read locus-level analogue of the short-read soft-clip pile-up signature used by MGEfinder (Durrant et al. 2020), applied at locus scale rather than at single-insertion-boundary scale and thresholded for ONT alignment characteristics.

## References

Amaral M, Devos DP. (2024) The neglected giants: Uncovering the prevalence and functional groups of huge proteins in proteomes. PLOS Computational Biology 20:e1012459. doi:10.1371/journal.pcbi.1012459

Aramaki T, Blanc-Mathieu R, Endo H, Ohkubo K, Kanehisa M, Goto S, Ogata H. (2020) KofamKOALA: KEGG Ortholog assignment based on profile HMM and adaptive score threshold. Bioinformatics 36:2251–2252. doi:10.1093/bioinformatics/btz859

Chaumeil PA, Mussig AJ, Hugenholtz P, Parks DH. (2022) GTDB-Tk v2: memory friendly classification with the Genome Taxonomy Database. Bioinformatics 38:5315–5316. doi:10.1093/bioinformatics/btac672

Chklovski A, Parks DH, Woodcroft BJ, Tyson GW. (2023) CheckM2: a rapid, scalable and accurate tool for assessing microbial genome quality using machine learning. Nature Methods 20:1203–1212. doi: 10.1038/s41592-023-01940-w

Coenye T, Vandamme P. (2005) Organisation of the S10, spc and alpha ribosomal protein gene clusters in prokaryotic genomes. FEMS Microbiology Letters 242:117–126. doi:10.1016/j.femsle.2004.10.050

Durrant MG, Li MM, Siranosian BA, Montgomery SB, Bhatt AS. (2020) A bioinformatic analysis of integrative mobile genetic elements highlights their role in bacterial adaptation. Cell Host & Microbe 27:140–153.e9. doi:10.1016/j.chom.2019.10.022

Emms DM, Kelly S. (2019) OrthoFinder: phylogenetic orthology inference for comparative genomics. Genome Biology 20:238. doi:10.1186/s13059-019-1832-y

Glöckner J, Kube M, Shrestha PM, Weber M, Glöckner FO, Reinhardt R, Liesack W. (2010) Phylogenetic diversity and metagenomics of candidate division OP3. Environmental Microbiology 12:1218–1229. doi:10.1111/j.1462-2920.2010.02164.x

Hallgren J, Tsirigos KD, Pedersen MD, Almagro Armenteros JJ, Marcatili P, Nielsen H, Krogh A, Winther O. (2022) DeepTMHMM predicts alpha and beta transmembrane proteins using deep neural networks. bioRxiv 2022.04.08.487609. doi:10.1101/2022.04.08.487609

Harshey RM. (2014) Transposable phage Mu. Microbiology Spectrum 2:MDNA3-0007-2014. doi: 10.1128/microbiolspec.MDNA3-0007-2014

Hyatt D, Chen GL, LoCascio PF, Land ML, Larimer FW, Hauser LJ. (2010) Prodigal: prokaryotic gene recognition and translation initiation site identification. BMC Bioinformatics 11:119. doi: 10.1186/1471-2105-11-119

Jeske O, Schüler M, Schumann P, Schneider A, Boedeker C, Jogler M, Bollschweiler D, Rohde M, Mayer C, Engelhardt H, Spring S, Jogler C. (2015) Planctomycetes do possess a peptidoglycan cell wall. Nature Communications 6:7116. doi:10.1038/ncomms8116

Jones P, Binns D, Chang HY, Fraser M, Li W, McAnulla C, McWilliam H, Maslen J, Mitchell A, Nuka G, Pesseat S, Quinn AF, Sangrador-Vegas A, Scheremetjew M, Yong SY, Lopez R, Hunter S. (2014) InterProScan 5: genome-scale protein function classification. Bioinformatics 30:1236–1240. doi: 10.1093/bioinformatics/btu031

Kizina J, Jordan SFA, Martens GA, Lipp JS, Hinrichs KU, Spormann AM. (2022) Methanosaeta and “Candidatus Velamenicoccus archaeovorus”. Applied and Environmental Microbiology 88:e02407–21. doi:10.1128/aem.02407-21

Larralde M. (2022) Pyrodigal: Python bindings and interface to Prodigal, an efficient method for gene prediction in prokaryotes. Journal of Open Source Software 7:4296. doi:10.21105/joss.04296

Nielsen TN. (2026a) Cohort-HMM marker recruitment with per-OG orthology QC for phylogenomic supermatrices. bioRxiv 2026.05.27.728348. doi:10.64898/2026.05.27.728348

Nielsen TN. (2026b) Omnitrophota dominate per-genome cross-replicon lateral gene transfer in a Fennoscandian deep-groundwater metagenome. bioRxiv 2026.05.20.726742. doi: 10.64898/2026.05.20.726742

Perez-Molphe-Montoya E, Küsel K, Overholt WA. (2022) Redefining the phylogenetic and metabolic diversity of phylum Omnitrophota. Environmental Microbiology 24:5437–5449. doi: 10.1111/1462-2920.16170

Reva ON, Tümmler B. (2008) Think big — giant genes in bacteria. Environmental Microbiology 10:768–777. doi:10.1111/j.1462-2920.2007.01500.x

Rinke C, Schwientek P, Sczyrba A, Ivanova NN, Anderson IJ, Cheng JF, Darling A, Malfatti S, Swan BK, Gies EA, Dodsworth JA, Hedlund BP, Tsiamis G, Sievert SM, Liu WT, Eisen JA, Hallam SJ, Kyrpides NC, Stepanauskas R, Rubin EM, Hugenholtz P, Woyke T. (2013) Insights into the phylogeny and coding potential of microbial dark matter. Nature 499:431–437. doi:10.1038/nature12352

Seymour CO, Palmer M, Becraft ED, Stepanauskas R, Friel AD, Schulz F, Woyke T, Hedlund BP, et al. (2023) Hyperactive nanobacteria with host-dependent traits pervade Omnitrophota. Nature Microbiology 8:727–744. doi:10.1038/s41564-022-01319-1

Shaw J, Marin MG, Li H. (2026) High-resolution metagenome assembly for modern long reads with myloasm. Nature Biotechnology (advance online publication). doi:10.1038/s41587-026-03053-z

West-Roberts J, Valentin-Alvarado L, Mullen S, Sachdeva R, Smith J, Hug LA, Gregoire DS, Liu W, Lin TY, Husain G, Amano Y, Ly L, Banfield JF. (2023) Giant genes are rare but implicated in cell wall degradation by predatory bacteria. bioRxiv 2023.11.21.568195. doi:10.1101/2023.11.21.568195

Zhang M, et al. (2023) Unexplored diversity and ecological functions of transposable phages. The ISME Journal 17:1015–1028. doi:10.1038/s41396-023-01414-z

