## Supplementary material for "A complete-genome view of phylum Omnitrophota and a multi-order capacity for very long proteins": Supplementary_Information.pdf

---

#### Supplementary Figures

**Figure S1 — Per-order metabolic profile.** KofamScan-derived KO presence/absence by order, validated against the cross-check.

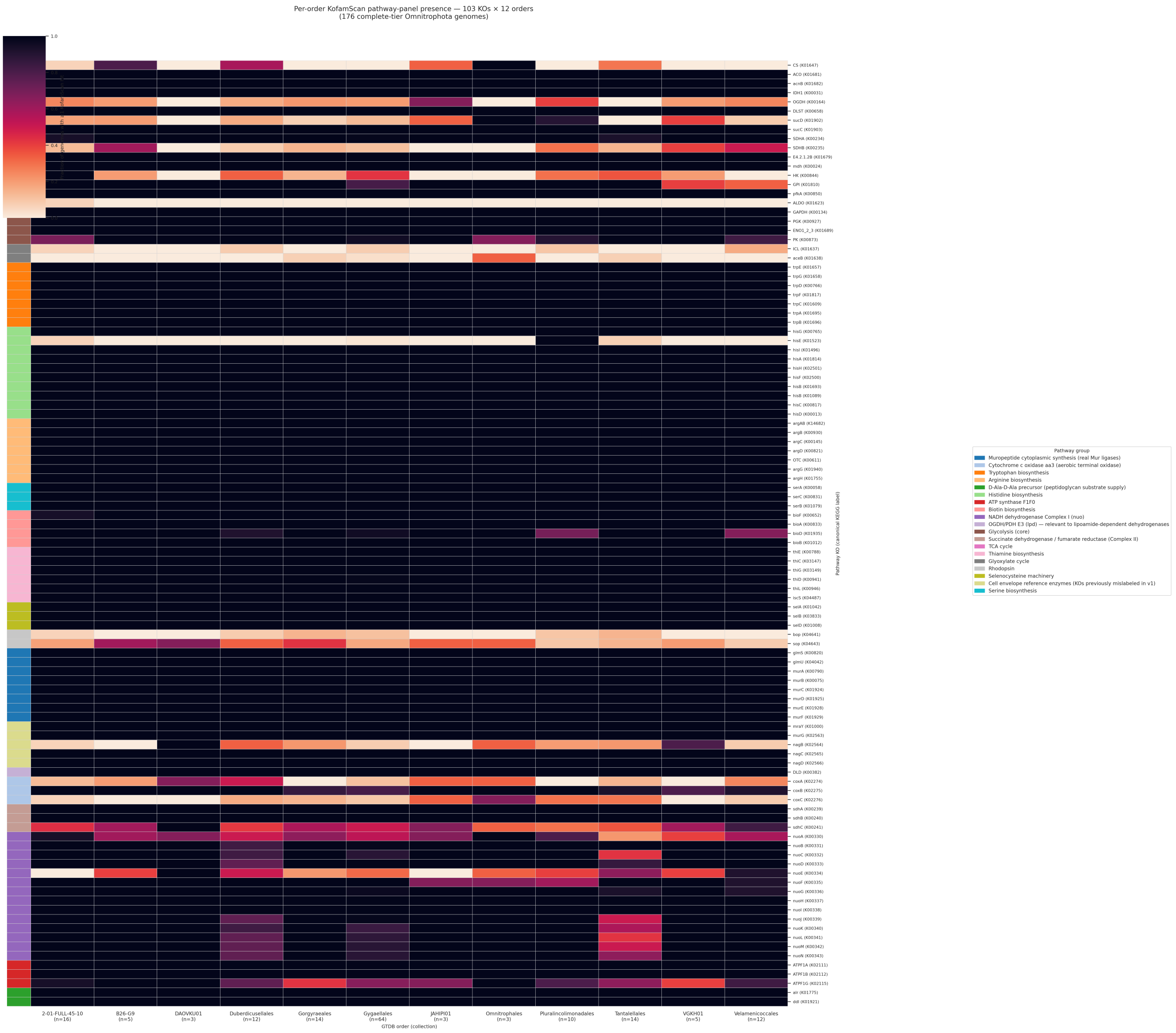

**Figure S2 — Extended 938-tip phylogeny.** Same 97-SC-OG marker set extended to all GTDB R232 *Omnitrophota* species representatives via the cohort-HMM recruitment pipeline + per-OG gene-tree QC + per-MAG paralog-rate filter (Nielsen 2026a). 229 in-house cohort + 4 PVC outgroups + 705 NCBI species representatives. Provides taxonomic placement of the new genomes against the global MAG inventory; not load-bearing for any main-text claim.

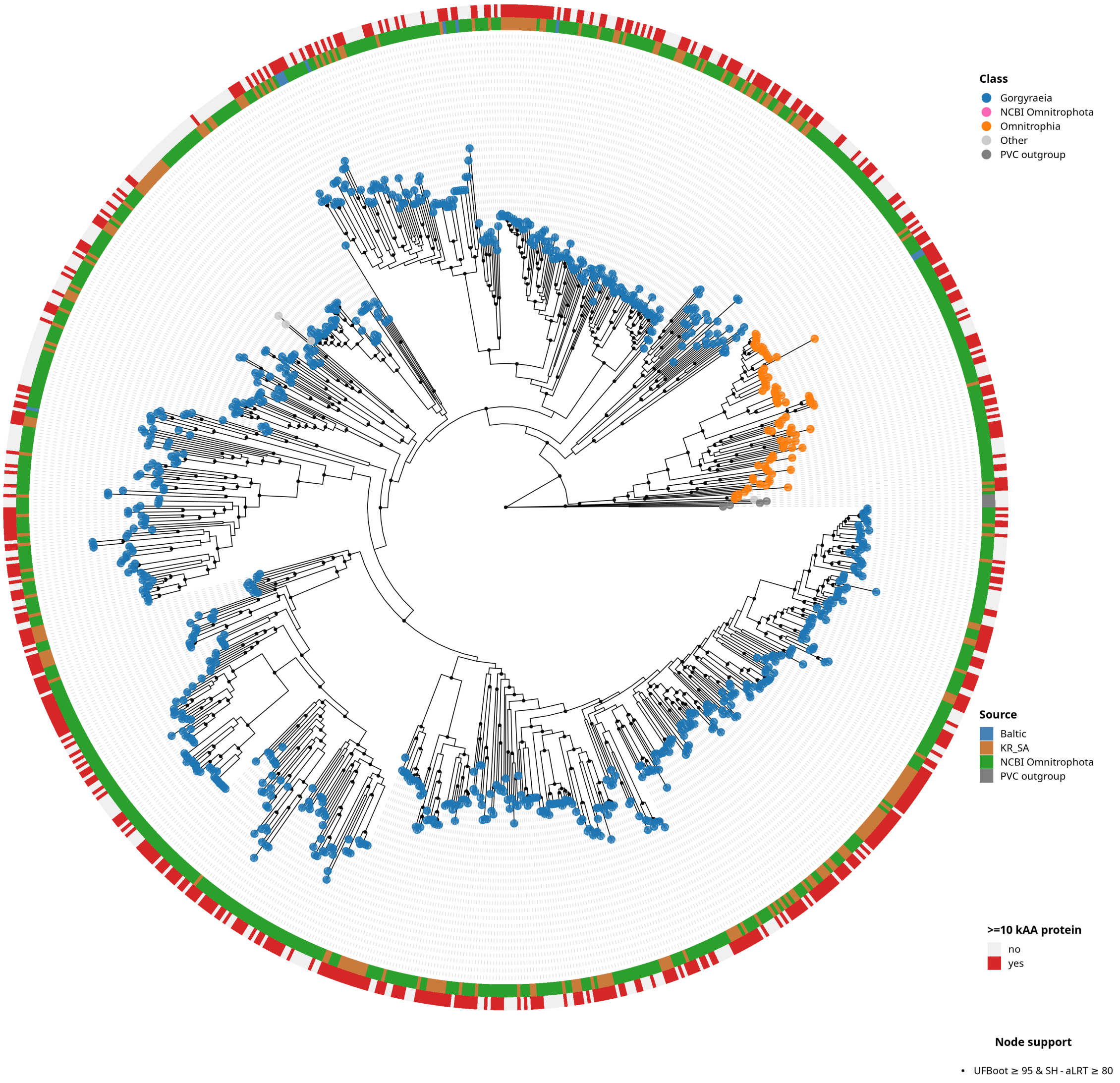

**Figure S3 — Assembly contiguity governs long-protein recovery.** Per-genome long-protein count by tier (panel A) and contig-count dependence within the 714 NCBI HQ *Omnitrophota* MAG population (panel B). Panel A: 93.8 % of the 176 complete genomes carry at least one ≥ 10-kAA protein, vs 81.1 % of the 53 in-house HQ MAGs at the same CheckM2 threshold and 42.9 % of 714 NCBI HQ MAGs. Panel B: Spearman $\rho = -0.36$ ($p < 1 \times 10^{-23}$) between contig count and long-protein count across the 714 NCBI MAGs; the per-contig-bin fraction with ≥ 1 long protein falls monotonically from 100 % (1–5 contigs) to 17.9 % (201–500 contigs). The standard CheckM2-HQ threshold passes MAGs whose fragmentation systematically truncates ≥ 10-kAA ORFs at contig boundaries, masking the long-protein landscape that the complete tier resolves. Both effects survive stratification by GTDB-Tk R232 order (Table S2): the complete vs NCBI tier gap is positive in all 11 orders with ≥ 3 representatives in both tiers, and the within-NCBI contig-count correlation is negative in all 9 orders with n ≥ 20 (significant at $p < 0.05$ in 8 of 9). Phylogenetic composition is therefore not a confounder for either signal. The 11 of 176 complete-tier collection genomes that themselves lack a ≥ 10 kAA protein cluster in just two orders — 8 in *Pluralincolimonadales* (a structurally distinct order whose long-protein-poor phenotype is consistent with its order-specific metabolic specializations) and 3 in *B26-G9* (all near-miss with longest predicted protein 7.8–9.6 kAA, just under the threshold; Table S2c).

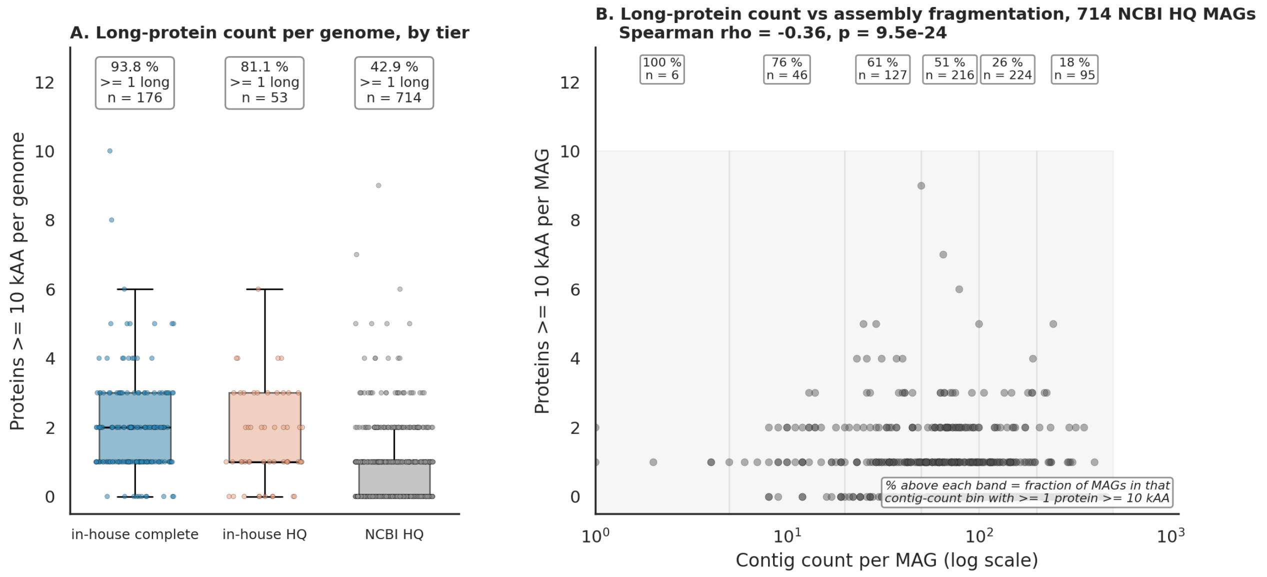

### Supplementary Tables

**Table S1 — Per-order/family breakdown of the 229 collection genomes.** GTDB-Tk R232 classifications; complete-tier and high-quality-tier counts shown separately. Sorted by class, then by complete count. Generated by `scripts/58_supplementary_table_S1.py` from `logs/01_cohort_manifest.tsv`.

| Class | Order | Family | Complete | High quality | Total |
|---|---|---|---|---|---|
| *Gorgyraeia* | *Gygaellales* | *Profunditerraquicolaceae* | 63 | 14 | 77 |
| *Gorgyraeia* | *2-01-FULL-45-10* | *2-01-FULL-45-10* | 16 | 3 | 19 |
| *Gorgyraeia* | *Gorgyraeales* | *Gorgyraeaceae* | 14 | 5 | 19 |
| *Gorgyraeia* | *Tantalellales* | *Tantalellaceae* | 14 | 4 | 18 |
| *Gorgyraeia* | *Duberdicusellales* | *4484-171* | 9 | 2 | 11 |
| *Gorgyraeia* | *Velamenicoccales* | *JABMQK01* | 8 | 3 | 11 |
| *Gorgyraeia* | *Pluralincolimonadales* | *Pluralincolimonadaceae* | 7 | 7 | 14 |
| *Gorgyraeia* | *B26-G9* | *B26-G9* | 5 | 2 | 7 |
| *Gorgyraeia* | *Velamenicoccales* | *Velamenicoccaceae* | 4 | 4 | 8 |
| *Gorgyraeia* | *Duberdicusellales* | *Duberdicusellaceae* | 3 | 1 | 4 |
| *Gorgyraeia* | *VGKH01* | *VGKH01* | 3 | 1 | 4 |
| *Gorgyraeia* | *DAOVKU01* | *DAOVKU01* | 3 | 0 | 3 |
| *Gorgyraeia* | *JAHIPI01* | *JAHIPI01* | 3 | 0 | 3 |
| *Gorgyraeia* | *Pluralincolimonadales* | *CAZUZR01* | 2 | 1 | 3 |
| *Gorgyraeia* | *(unnamed)* | *(unnamed)* | 2 | 0 | 2 |
| *Gorgyraeia* | *JABMRJ01* | *JABMRJ01* | 2 | 0 | 2 |
| *Gorgyraeia* | *VGKH01* | *(unnamed)* | 2 | 0 | 2 |
| *Gorgyraeia* | *JAFGHF01* | *JAFGHF01* | 1 | 1 | 2 |
| *Gorgyraeia* | *B26-G9* | *JBDRCZ01* | 1 | 0 | 1 |
| *Gorgyraeia* | *Gygaellales* | *JAHJCI01* | 1 | 0 | 1 |
| *Gorgyraeia* | *JABMRG01* | *JABMRG01* | 1 | 0 | 1 |
| *Gorgyraeia* | *JABMRG01* | *JAFGKX01* | 1 | 0 | 1 |
| *Gorgyraeia* | *JABMRN01* | *JABMRN01* | 1 | 0 | 1 |
| *Gorgyraeia* | *JAHJTE01* | *JAHJTE01* | 1 | 0 | 1 |
| *Gorgyraeia* | *JAZDFF01* | *JAZDFF01* | 1 | 0 | 1 |
| *Gorgyraeia* | *JBDRLY01* | *JBDRLY01* | 1 | 0 | 1 |
| *Gorgyraeia* | *VGKN01* | *VGKN01* | 1 | 0 | 1 |
| *Gorgyraeia* | *Zapsychrales* | *JANLHA01* | 1 | 0 | 1 |
| *Gorgyraeia* | *JABMRN01* | *RBG-13-46-9* | 0 | 1 | 1 |
| *Gorgyraeia* | *JAUYCX01* | *JBMONJ01* | 0 | 1 | 1 |
| *Gorgyraeia* | *Zapsychrales* | *JBDRMG01* | 0 | 1 | 1 |

| Class | Order | Family | Complete | High quality | Total |
|---|---|---|---|---|---|
| *Omnitrophia* | *Omnitrophales* | *Aquincolibacteriaceae* | 3 | 2 | 5 |
| *Omnitrophia* | *JAHJGZ01* | *JAHJGZ01* | 2 | 0 | 2 |
| **Total** | | | **176** | **53** | **229** |

**Table S2 — Per-order stratification of the long-protein-by-tier effect** (companion to Figure S3). Demonstrates that the in-house complete vs NCBI HQ gap and the within-NCBI contig-count effect both persist after stratifying by GTDB-Tk R232 order — phylogenetic composition is not a confounder. Generated by `scripts/60_supplementary_table_S2.py` from `logs/01_cohort_manifest.tsv`, `logs/long_proteins_cross_sample_ge10kaa.tsv`, `logs/long_proteins_ncbi_ge10kaa.tsv`, `ncbi/02_ncbi_hq_candidates.tsv`, and the GTDB R232 `bac120_metadata_r232.tsv`.

#### Table S2a — Per-order tier gap (orders with ≥ 3 representatives in both in-house complete and NCBI HQ)

| Order | n complete | n HQ | n NCBI | complete % | HQ % | NCBI % | gap (complete − NCBI, pp) |
|---|---|---|---|---|---|---|---|
| *JAHIPI01* | 3 | - | 4 | 100.0 | - | 0.0 | +100.0 |
| *Tantalellales* | 14 | 4 | 49 | 100.0 | 75.0 | 30.6 | +69.4 |
| *VGKH01* | 5 | 1 | 3 | 100.0 | 100.0 | 33.3 | +66.7 |
| *Gorgyraeales* | 14 | 5 | 25 | 100.0 | 80.0 | 40.0 | +60.0 |
| *Duberdicusellales* | 12 | 3 | 59 | 100.0 | 100.0 | 40.7 | +59.3 |
| *2-01-FULL-45-10* | 16 | 3 | 28 | 100.0 | 100.0 | 42.9 | +57.1 |
| *Gygaellales* | 64 | 14 | 165 | 100.0 | 100.0 | 43.6 | +56.4 |
| *Omnitrophales* | 3 | 2 | 78 | 100.0 | 100.0 | 55.1 | +44.9 |
| *B26-G9* | 6 | 2 | 15 | 50.0 | 0.0 | 13.3 | +36.7 |
| *Velamenicoccales* | 12 | 7 | 20 | 100.0 | 100.0 | 65.0 | +35.0 |
| *Pluralincolimonadales* | 9 | 8 | 16 | 11.1 | 25.0 | 0.0 | +11.1 |

The complete-vs-NCBI gap is positive in all 11 orders, with median 56.4 percentage points. *Pluralincolimonadales* is the lone order where complete-tier rate is itself low (1 of 9 collection genomes carries a ≥ 10-kAA protein); even there the complete > NCBI ordering holds.

#### Table S2b — Within-NCBI contig-count effect (orders with n ≥ 20)

| Order | n | Spearman ρ | p |
|---|---|---|---|
| *2-02-FULL-51-18* | 20 | −0.693 | $7.0 \times 10^{-4}$ |
| *Tantalellales* | 49 | −0.528 | $9.5 \times 10^{-5}$ |
| *Velamenicoccales* | 20 | −0.501 | $2.4 \times 10^{-2}$ |
| *Omnitrophales* | 78 | −0.491 | $5.1 \times 10^{-6}$ |
| *Gorgyraeales* | 25 | −0.478 | $1.6 \times 10^{-2}$ |

| Order | n | Spearman ρ | p |
|---|---|---|---|
| *Gygaellales* | 165 | −0.467 | $2.6 \times 10^{-10}$ |
| *Zapsychrales* | 125 | −0.333 | $1.5 \times 10^{-4}$ |
| *Duberdicusellales* | 59 | −0.312 | $1.6 \times 10^{-2}$ |
| *2-01-FULL-45-10* | 28 | −0.187 | $3.4 \times 10^{-1}$ |

The within-NCBI contig_count vs n_ge10k correlation is negative in all 9 orders, significant at $p < 0.05$ in 8 of 9. The lone non-significant order (*2-01-FULL-45-10*) is the smallest in this subset (n = 28) and is consistent in direction.

#### Table S2c — Per-order distribution of the 11 complete-tier collection genomes that lack a ≥ 10 kAA protein

##### Order-level summary:

| Order | n in collection | n missing | % missing |
|---|---|---|---|
| *Pluralincolimonadales* | 9 | 8 | 88.9 |
| *B26-G9* | 6 | 3 | 50.0 |

Every other order in the 176-genome complete tier has 0 missing genomes. The 11 misses are exhaustively from these two orders.

##### Per-genome detail:

| Genome | Order | Longest predicted protein (AA) |
|---|---|---|
| SA1420A_u25336563 | *B26-G9* | 9,625 |
| SA1420A_u14286630 | *B26-G9* | 9,526 |
| SA1420A_u15005499 | *B26-G9* | 7,803 |
| KR0015B_u28424469 | *Pluralincolimonadales* | 3,208 |
| KR0015B_u18943763 | *Pluralincolimonadales* | 3,119 |
| KR0015B_u15048077 | *Pluralincolimonadales* | 2,531 |
| KR0015B_u740188 | *Pluralincolimonadales* | 2,435 |
| KR0015B_u1846880 | *Pluralincolimonadales* | 2,333 |
| KR0015B_u18481865 | *Pluralincolimonadales* | 2,187 |
| KR0015B_u31766398 | *Pluralincolimonadales* | 2,016 |
| SA1420A_u19858292 | *Pluralincolimonadales* | 1,980 |

Two different mechanisms read off this table. *Pluralincolimonadales* (8/11) is a genuinely long-protein-poor lineage — longest predicted proteins span 2.0–3.2 kAA, far below the ≥ 10 kAA threshold and well below any reasonable "near-miss" margin. This is consistent with the order's distinct metabolic specializations reported elsewhere in this paper (Discussion §"A constrained metabolic profile…"). *B26-G9* (3/11) is a near-miss case: all three missing genomes carry their longest predicted protein at 7.8–9.6 kAA — just under the 10 kAA cutoff — while the other 3 of 6 *B26-G9* collection genomes do carry at least one ≥ 10 kAA protein. *B26-G9* therefore has the phenotype at a slightly smaller length scale; the 10 kAA threshold catches only the upper half of its distribution.
